# Learning a threat converges on the circuit processing innate threat

**DOI:** 10.64898/2026.08.26.747219

**Authors:** Noëmie Mermet-Joret, Milad Nazari, Andreas T. Pommer, Sanaz Ansarifar, Joana Silva Luz, Anne-Katrine Vestergaard, Sadegh Nabavi

## Abstract

A prevailing view in affective neuroscience holds that innate and learned behaviors are processed through distinct neuroanatomical pathways, one pre-wired, the other running on synaptic plasticity. However, here we show that processing innate and learned threats in the lateral amygdala deviates fundamentally from this view. We tracked the three core elements of circuit function—excitatory neurons, inhibitory neurons, and neuromodulators—in mice, as they were exposed to an innately aversive looming stimulus and as they learned a cued threat. Tracking the same neurons across sessions, revealed a subpopulation of excitatory neurons recruited by the innate threat that was preferentially potentiated following auditory threat learning. Furthermore, both forms of threat converged on the same modulatory mechanisms: the disinhibitory VIP–SST motif and norepinephrine release, but with a critical difference. While the innately aversive stimulus possessed privileged access to these pathways, the learned cue acquired access through synaptic plasticity. In this instance, learning about a new threat apparently recruits a circuit that protects animals from natural threats.

## Introduction

A dominant perspective in studies of fear and threat processing holds that innate and learned threats, though driving similar defensive behaviors, rely on distinct neurocircuitry^1,2^; one, pre-wired, is evolved through natural selection, whereas the other emerges through experience-dependent synaptic plasticity following prior exposure to threat signals. Supporting this view, an animal’s response to a natural predator— an innate threat— is processed within subcortical regions, such as the hypothalamic nuclei and medial amygdala^3–8^. On the other hand, learning about a threat— for example, associating a previously neutral stimulus with an aversive outcome— relies on cortical-like structures, such as the basolateral amygdala and the hippocampus ^9–19^.

This dichotomy, however, has been shown to have notable exceptions. The most striking example is the lateral part of the amygdala (LA), a region essential for almost any known form of threat learning ^9–13,20–22^. Lesioning the LA in rodents eliminates defensive responses to a live predator as well as to an expanding overhead stimulus, also known as a looming stimulus, mimicking an aerial predator^23–26^.

A looming stimulus is perhaps the most evolutionarily conserved form of innate threat^27^. It triggers rapid defensive responses in species as diverse as zebrafish, rodents and humans^27–32^. Notably, the same thalamic-amygdala pathway that implements plasticity for a learned aversive stimulus also processes the defensive response to a looming stimulus^25,26^. Put differently, a neutral tone and a looming stimulus both activate the LA, but only the loom triggers a defensive response on its own. The tone achieves this only by means of plasticity, following its association with an aversive experience such as an electric shock.

This suggests that learning a threat, through plasticity, might recruit the circuit within the LA that processes an innate threat. To test this hypothesis, we examined the three core elements of circuit function—excitatory neurons, inhibitory neurons, and neuromodulators— in mice as they were exposed to a looming stimulus signaling an innate threat and as they learned a new threat.

## Results

To monitor the activity of individual neurons across days, we performed microendoscopic calcium imaging of CaMKII-GCaMP8m–expressing amygdala principal neurons in freely behaving mice^33–36^. We confined our recordings to the lateral part of the amygdala (LA, Fig. 1A) as we have previously shown that the activity within this region is required for processing innate as well as learned aversive cues^26^.

**Figure 1.**
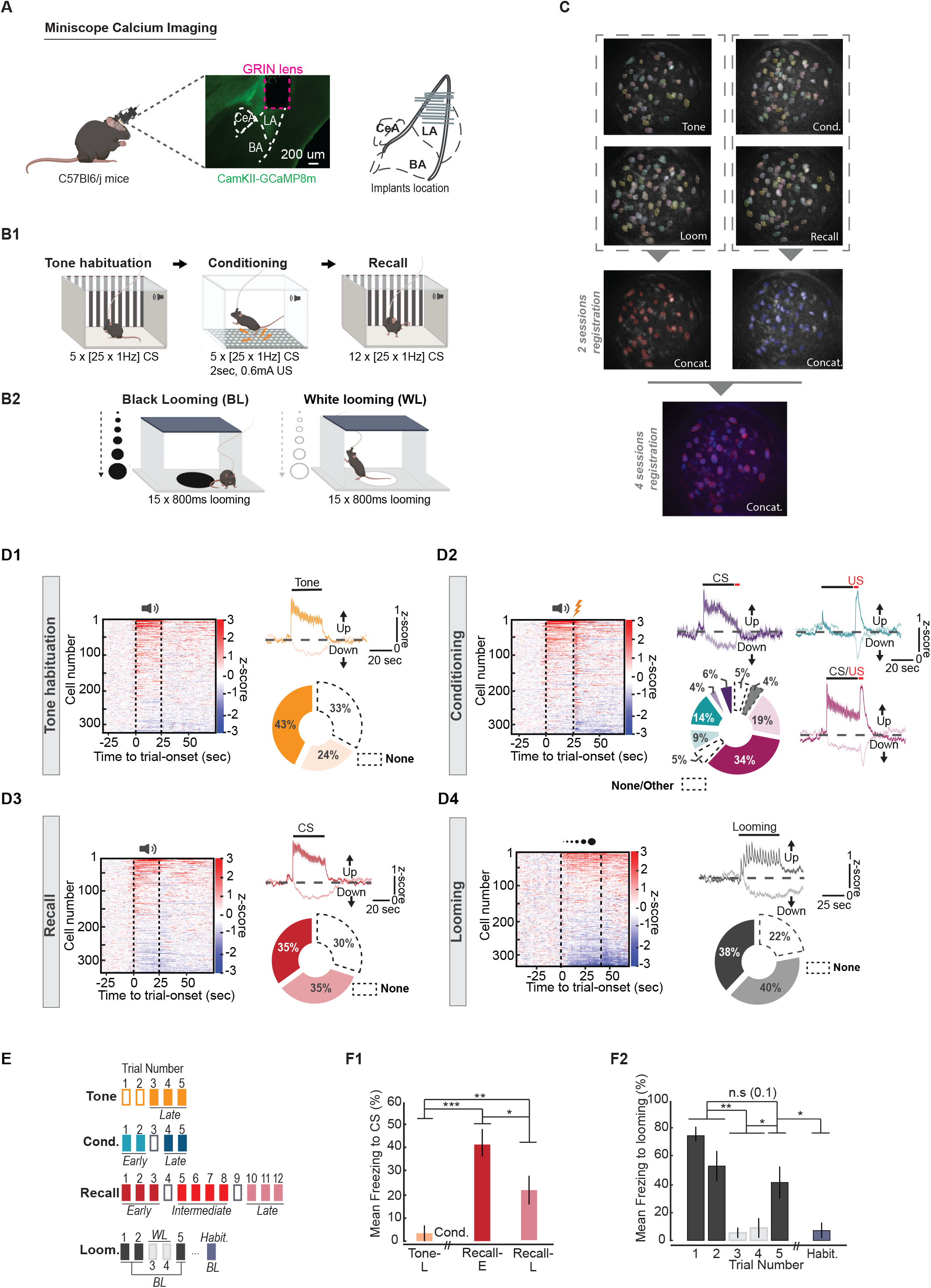
Longitudinal single-cell calcium imaging of the lateral amygdala principal neurons during exposure to learned and innately aversive cues. (A) Left, miniscope calcium imaging of CaMKII-GCaMP8m-expressing principal neurons in the lateral amygdala (LA) of freely behaving wild-type mice, with representative GCaMP8m expression and GRIN lens placement. Right, GRIN lens placement within the LA across all recorded animals. CeA, central amygdala; BA, basal amygdala. (B) Four-day behavioral paradigm for the tone habituation, auditory fear conditioning, and recall (B1), and black or white looming object exposure (B2). (C) Field-of-view and cross-session registration strategy of the same neurons across the Tone, Conditioning, Looming and Recall sessions. The colored region-of-interests (ROIs) represent individual neurons and are aligned across sessions. (D) Single-cell activity profiles for each session: tone habituation (D1), conditioning (D2), recall (D3), and looming (D4). Heatmaps show z-scored single-cell activity, sorted by their response amplitude from stimulus onset. Traces show mean (± s.e.m., shaded) z-scored activity averaged across trials for cells classified as activated (up) or suppressed (down). Donut charts show the proportion of activated, suppressed, and non-responsive (none) neurons for each session; for conditioning (D2), neurons are further classified by their response to the conditioned stimulus (CS), the unconditioned stimulus (US), or both (CS/US). The category Other includes neurons that showed opposite responses to CS and US, but were not further investigated. A neuron was classified as stimulus-responsive if its activity deviated significantly from baseline (Wilcoxon signed-rank test, P < 0.05). (E) Segmentation of each session into early (E), intermediate (I) and late (L) trial epochs. (F) Mean freezing response (F1) over 40 seconds following tone onset during Tone-L, Recall-E, and Recall-L (n=11 for Tone, n= 12 mice for Recall; Overall effect: Kruskal–Wallis test, χ^2^ = 19.02, d.f. = 2, *P* = 7.41 × 10^−5^. Pairwise comparisons: two-sided Mann–Whitney *U* tests. Tone vs Recall-E, *P* = 1.72 × 10^−4^; Tone vs Recall-L, *P* = 1.25 × 10^−3^; Recall-E vs Recall-L, *P* = 0.017. *P < 0.05, **P < 0.01, ***P < 0.001. Mean freezing response (F2) over 45 seconds following the onset of either black (BL) or white (WL) looming stimuli, and to BL habituation (Habit.) (n=9 mice; Friedman test, χ^2^ = 21.08, d.f. = 3, P = 1.02 × 10^−4^. Pairwise comparisons: two-sided Wilcoxon signed-rank tests. BL[1,2] vs WL[3,4], P = 0.0039; BL[1,2] vs BL[5], P = 0.1; WL[3,4] vs BL[5], P = 0.016; BL[5] vs BL[Habit.], P = 0.016. *P < 0.05, **P < 0.01; ns, not significant. The error-bars represent s.e.m.

The mice underwent a four-day experimental protocol (Fig. 1B), starting with the Tone day, followed by Loom, Conditioning, and Recall, an order chosen to prevent carryover from the more aversive sessions into the less aversive ones. For our learning threat protocol, first, we exposed the animals to a neutral pulsated tone (25 pulses (200 ms each) delivered at 1 Hz) in context A. To produce a learned aversive memory, we paired this tone (conditioned stimulus [CS]) with an aversive footshock (unconditioned stimulus [US]) in context B^35^. The following day, the mice, upon re-exposure to the CS in context A, showed freezing responses indicating successful learning of the association^37,38^.

As for the innately aversive threat, we used the looming stimulus, an overhead expanding black shadow that is thought to mimic an approaching aerial predator (Fig. 1B2). Unlike the auditory tone used as a learned threat, the looming shadow triggers defensive responses without prior learning^27,39^. As a non-aversive control, we used a white expanding shadow which does not elicit a defensive response ^27^.

We recorded 592 ± 35 cells in total across 12 mice per recording session. Using image alignment, we tracked the same neurons across the four sessions (Fig. 1C). A high percentage of tracked cells remained active across two consecutive sessions (72%), but this number dropped to 59% and 54% when tracking cells across three and four consecutive sessions. For analyses involving the black-looming shadow, we excluded three of the mice because of their weak defensive response to the stimulus.

We deemed a neuron to be encoding a given stimulus if its activity significantly deviated from the baseline (pValue < 0.05, Wilcoxon signed-rank test). For each stimulus, a neuron could show activation, suppression, or no significant change in response (Fig. 1D). Additionally, some neurons could respond to different stimuli during the same session. For example, during conditioning, 34% of all cells were activated by the tone and US (Fig. 1D2). Similarly, 19% of the neurons were inhibited by both the tone and US (Fig. 1D2). Over different days, a neuron could also be activated by one stimulus but suppressed by another (more on this later).

Informed by the behavioral observations, we separated the trials into early (E), intermediate (I) or late (L) trials for all the analyses presented here unless otherwise specified (Fig. 1E). For the ‘Tone’ day, we considered the last three trials (Tone-L), when the animals were habituated to the auditory cue; for the ‘Conditioning’ day, we distinguished the first two trials of conditioning (Cond-E) as an early stage of training from the last two trials of conditioning (Cond-L), where the animals effectively learned the association between CS and US. Similarly, for the ‘Recall’ day, we distinguished the first three presentations of the CS (Recall-E) where animals showed the highest percentage of freezing, from middle trials (Recall-I) and the last three trials where the animals started to show extinction to the cue (Recall-L) (Fig. 1F1). As for the ‘Looming’ day, we only considered the first three presentations of the black-looming (BL) stimulus where the animals showed the highest defensive response (Fig. 1F2).

### Learning reorganizes LA population dynamics through a distinct subset of tone-responsive neurons

To investigate how learning shapes population dynamics, we analyzed the LA population responses to the auditory tone before conditioning, when it was neutral, and after conditioning, when it evoked robust freezing (Fig. 2A; n = 12 mice). First, we quantified the functional similarity between neurons by computing pairwise cosine-similarity across trial-averaged activity traces for all neuron pairs (n = 8,498 pairs), separately for the Tone-L and Recall-E sessions. We observed a significant increase in the similarity coefficient following conditioning, consistent with a learning-related reorganization in which neurons developed increasingly similar tone-response profiles (Fig. 2B).

**Figure 2.**
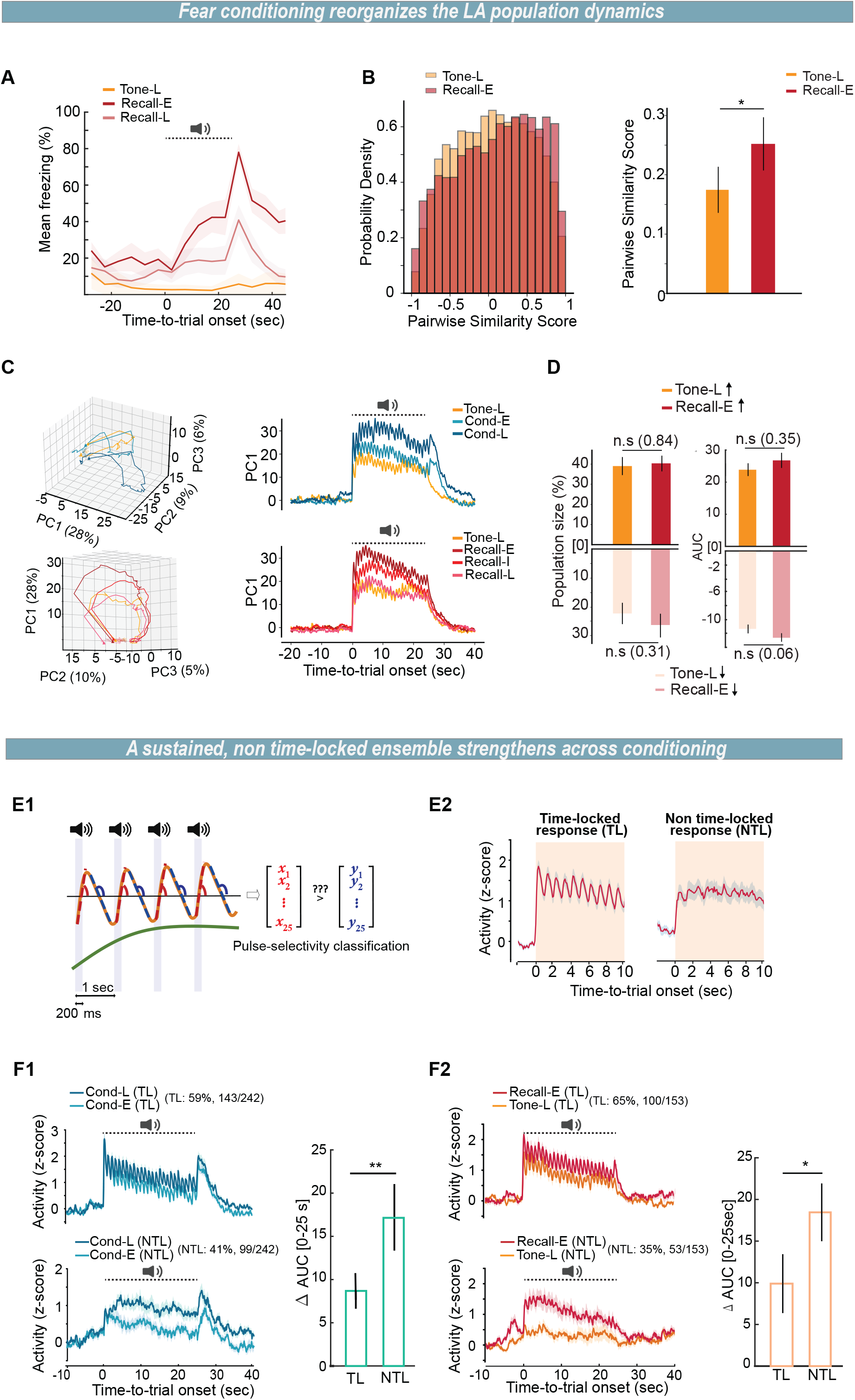
Fear conditioning reorganizes LA population dynamics and is accompanied by an enhanced response in a population ensemble with sustained, non-time-locked activity. (A) Freezing percentage to the CS (dashed line) over time for Tone-L, Recall-E, and Recall-L. Shaded areas represent s.e.m. (B) Left, Distribution of pairwise cosine-similarity scores computed across trial-averaged activity for all neuron pairs (n=8,498 pairs from n=12 mice), for Tone-L versus Recall-E; Right, The averaged similarity score increases after conditioning (Tone-L, 0.17 ± 0.04; Recall-E, 0.25 ± 0.04; per mouse median, paired t-test, p = 0.02; error bars represent s.e.m.). (C) Principal Component Analysis; Left, population trajectories in PC1– PC3 state space (with % of variance) for the Tone vs Conditioning sessions (top) and the Tone vs Recall sessions (bottom); Right, amplitude of PC1 projection over time, during the CS period, across Tone-L, Cond-E and Cond-L (top) and across Tone-L, Recall-E, Recall-I and Recall-L (bottom). (D) Single-cell analysis comparing the fraction of tone-responsive neurons (Up/Down) and area under curve (AUC, Up/Down) for Tone-L versus Recall-E (paired t-test; n.s, non-significant, errorbars represent s.e.m.). (E) Classification of CS-activated neurons into two distinct populations based on their pulse-selectivity profile; (E1) Schematic of the pulse-selectivity classification distinguishing time-locked (TL) from non-time-locked (NTL) responses to individual CS pulses; (E2) Representative averaged activity for both TL and NTL populations. (F) CS-evoked responses of TL and NTL neurons across learning; (F1) Activity of TL (top) and NTL (bottom) neurons during early (Cond-E) and late (Cond-L) conditioning (n = 242 CS-activated neurons; TL, 59%, 143/242 neurons; NTL, 41%, 99/242 neurons), and difference in AUC (ΔAUC, 0–25 s) between early and late trials for both ensembles (Mann-Whitney U test, P = 0.005); (F2) Activity of TL (top) and NTL (bottom) neurons during Tone-L and Recall-E (n = 153 CS-activated neurons; TL, 65%, 100/153 neurons; NTL, 35%, 53/153 neurons), and ΔAUC (0–25 s) between the two sessions for both ensembles (Mann-Whitney U test, P = 0.016). Error bars represent s.e.m.

The pairwise similarity, however, provides only a static, session-by-session snapshot of the population. It cannot capture how the tone response evolved as learning proceeded. Therefore, we sought a common reference frame in which the population activity could be followed continuously within and across sessions. For this, we performed Principal Component Analysis (PCA) on the concatenated z-scored population activity across the Tone, Conditioning, and Recall sessions (Fig. 2C, Supplementary Fig. 1A). The first principal component (PC1) accounted for 28% of the variance across the three sessions, whereas the second and third components contributed about 10% and 6% (Supplementary Fig. 1A).

The PC1 projection peaked at tone onset and dropped after tone offset, indicating that PC1 captured the tone-evoked component of the population response. Importantly, the magnitude of this projection increased from late tone to late conditioning and early recall (Fig. 2C). Specifically, within the conditioning session, the projection grew as training progressed. In contrast, during recall, the initially high projection declined with successive CS presentations, ultimately returning to the level seen in the tone session.

Notably, this graded profile correlated with that of the animals’ defensive response, which was likewise strongest early and diminished by the late stage of recall (Fig. 2A), consistent with PC1 tracking the salience of the learned cue. Importantly, PC1 is not a readout of the behavior itself: it remained confined to the tone and returned to baseline while freezing was still elevated, and the tone alone (Tone-L) evoked a substantial PC1 response despite eliciting little freezing (Fig. 2A,C).

While functional similarity analysis and PCA indicate learning-induced plasticity in the LA, they do not provide insight into how the plasticity is implemented mechanistically. We therefore performed single-cell analysis, quantifying both the fraction of CS-responsive neurons and their response amplitude following training. This, however, did not reveal a significant change— neither in population size nor in AUC response— from Tone to Recall sessions (Fig. 2D). This population-level measure considered all tone-responsive neurons and could therefore mask a response increase confined to a subset of cells. We, therefore, probed for the neurons that increased their responses to the tone from Tone to Recall sessions, regardless of their selectivity on Tone day. This showed that among all CS-activated neurons on the Recall session, about 40% had at least doubled their response compared to the Tone session (Supplementary Fig. 1B).

Upon closer examination, we observed that these strongly potentiated neurons had distinct response dynamics from those of the non-potentiated CS-responsive neurons (Supplementary Fig. 1B; Fig. 2E). Specifically, they showed sustained activity throughout the pulsated tone period, while the rest of the CS-responsive neurons exhibited a tightly aligned response to each 200 ms auditory pulse with clear rises and falls. To validate this, we independently classified neurons based on their response dynamics — sustained (or non-time-locked, NTL) versus time-locked (TL) neurons (Fig. 2E) — and we then examined how their CS-evoked responses changed with conditioning and from tone to recall. This approach converged onto the same conclusion: neurons with sustained activity (NTL neurons) not only exhibited a significantly greater increase in CS-evoked responses as conditioning progressed (Fig. 2F1), but were also the population showing a more significant enhancement from Tone to Recall (Fig. 2F2).

### Aversive and non-aversive looming are encoded by distinct LA excitatory populations

To investigate how the LA encodes an innately aversive cue, we exposed the animals to multiple presentations of overhead black-looming stimulus (BL) while recording the excitatory neuronal activity. As a control for a visual stimulus without valence, we used a white looming stimulus (WL) within the same session^26,27^ (Fig. 3A, n = 9 mice). Approximately one third of the recorded neurons were activated, and another one-third were suppressed in response to the BL (Fig. 3B-D, Fig. 1D4). A considerable subset of neurons showed stable selectivity across the BL trials, with significant overlap between the neurons activated on separate presentations (BL[T1] to BL[T3]), indicating relatively stable recruitment of a defined ensemble. By contrast, the neurons activated by the BL were significantly less likely to be activated by the WL (Fig. 3D).

**Figure 3.**
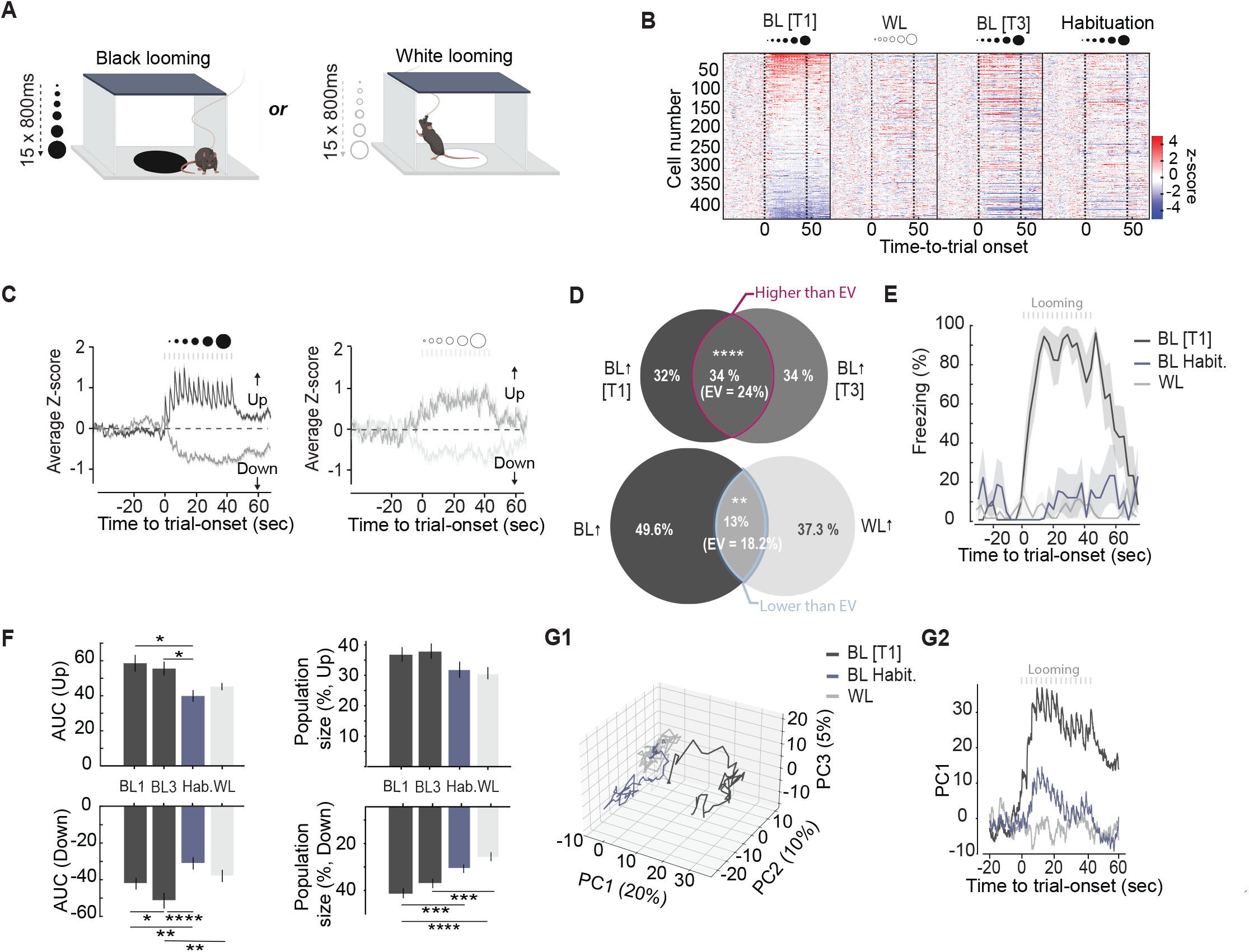
A subset of LA principal neurons is activated by the aversive looming stimulus. (A) Freely moving mice were presented with repeated trials of 15 looming stimuli (800 ms each), using either an aversive black disk (black looming, BL) or a non-aversive white disk (white looming, WL). (B) Heatmaps of single-cell z-scored activity across four conditions (BL[T1], WL, BL[T3], habituation) from 9 mice. Cells are ranked by their response amplitude on the first BL trial (BL[T1]). Dotted vertical lines indicate stimulus onset and offset. (C) Average z-scored population activity for BL (left) and WL (right) trials, showing activated (Up) and suppressed (Down) responses. Shaded areas represent s.e.m.; gray bars indicate individual looming stimuli. (D) Venn diagrams showing overlap between LA neurons significantly activated by BL or WL across trials. (Top) Overlap between BL[T1]- and BL[T3]-responsive neurons (34% shared; expected overlap by chance [EV] = 24%; P < 0.0001), indicating a stable population reactivated across repeated BL presentations. (Bottom) Overlap between BL- and WL-responsive neurons (all trials, 13% shared; EV = 18.2%; P < 0.01), indicating that neurons activated by BL and WL largely belong to distinct populations. P-values were calculated using a two-sided Fisher’s exact test. (E) Freezing behavior (%) over time for BL[T1], BL habituation, and WL trials. Black, but not white, looming elicits robust freezing, which declines with habituation. Gray vertical bars indicate individual looming stimuli. Shading represents s.e.m. n= 9 mice. (F) Quantification of population responses across conditions (BL[T1], BL[T3], Hab., WL). (Left) Per-cell area under the curve (AUC) for activated (top, ‘Up’, one-way ANOVA: p = 0.0038, followed by post hoc Tukey-Kramer) and suppressed (bottom, ‘Down’, one-way ANOVA: p < 0.0001, followed by post hoc Tukey-Kramer) populations; (Right) Proportion of cells classified as activated (top, ‘Up’, one-way ANOVA: p = 0.0462, Post-hoc Tukey–Kramer comparisons did not identify any single trial pair as significant after correction for multiple comparisons) or suppressed (bottom, ‘Down’, one-way ANOVA: p < 0.0001, followed by post hoc Tukey-Kramer). p-values * < 0.05, ** < 0.01, *** < 0.001, **** < 0.0001. Error bars represent s.e.m. (G) Principal component analysis (PCA) of LA population activity (PC1, 20% variance; PC2, 10%; PC3, 5%). (G1) Three-dimensional state-space trajectory (PC1–PC3) for BL[T1], BL habituation, and WL, illustrating separable population-level representations. (G2) PC1 projection over time for the same three conditions.

As the session progressed, the animals showed a rapid habituation to the BL, with no visible defensive response after several trials of BL exposure (Fig. 3E). The activity of the neurons in the LA reflected this behavioral change. The absolute amplitude of the response for the neurons activated or suppressed by the BL declined significantly during the final trials when defensive responding was at its minimum (Fig. 3F). Meanwhile, the proportion of the neurons suppressed by the BL decreased over trials, while the number of BL-activated neurons remained comparable between the early and later trials (Fig. 3F). Overall, these results suggest that both BL-activated and BL-suppressed LA neurons contribute to the expression of innate defensive behavior.

In the previous section, we showed that the magnitude of the first principal component (PC1) projection tracked the salience of the learned aversive cue rather than the defensive response it evoked (Fig. 2A,C). To test whether the same holds for an innately aversive cue, we performed a PCA on the z-scored population activity (Fig. 3G). In the three-dimensional state space defined by PC1–PC3, trajectories corresponding to the first BL trial, the BL habituation trial, and non-aversive WL exposure occupied clearly distinct regions (Fig. 3G1). The first BL trial was separated from both BL habituation and WL, which in turn lay closer to each other than either did to the first BL trial. These results support the notion that aversive and non-aversive visual stimuli are represented by distinct patterns of population activity within the LA.

The first principal component accounted for 20% of the variance, whereas the second and third explained substantially less (∼10% and ∼5%). During the first BL trial, the magnitude of the PC1 projection rose following stimulus onset and remained elevated beyond stimulus offset (Fig. 3G2), accompanied by robust freezing that likewise persisted beyond offset (Fig. 3E). This sustained projection contrasted sharply with the BL habituation trial, where PC1 was markedly reduced and no longer showed a prolonged post-stimulus increase. Projections during WL were minimal and remained near baseline. Notably, PC1 during the habituation trial remained elevated despite freezing having returned to near-baseline levels (Fig. 3E), suggesting that the population response does not simply track the expression of defensive behavior.

### Excitatory neurons activated by the innate threat are suppressed by the US, but recruited by the learned threat

At this stage, our data show that excitatory neurons in the LA encode both learned and innate aversive stimuli. Given only one requires plasticity, it may be concluded that they are implemented by distinct neuroanatomical structures and neurophysiological mechanisms. However, the opposite is also possible, given both are processed in the LA and produce similar defensive responses. To explore these possibilities, we began with an unsupervised, discovery-driven analysis which does not impose a specific hypothesis on the data. For this purpose, we first reduced the dimensionality of the data using PCA, taking each neuron’s trial-averaged response profile as input, so that every neuron was positioned as a single point in a three-dimensional PC space. (see Methods). Subsequently, we clustered the neurons in this space using the k-means algorithm.

Comparing the clusters between Conditioning and Loom sessions led to an unexpected observation. Given that both US and the BL stimulus are aversive, one would expect overlap between the neural clusters modulated in the same direction by these stimuli. However, we observed that the cluster representing the neurons activated during exposure to the BL significantly overlapped with the cluster representing the neurons suppressed by the US. At the same time, the clusters representing BL-activated and US-activated neurons overlapped less than expected by chance (Fig. 4B). The reverse was also true: the cluster for BL-suppressed neurons significantly overlapped with the cluster representing US-activated neurons (Fig. 4B).

**Figure 4.**
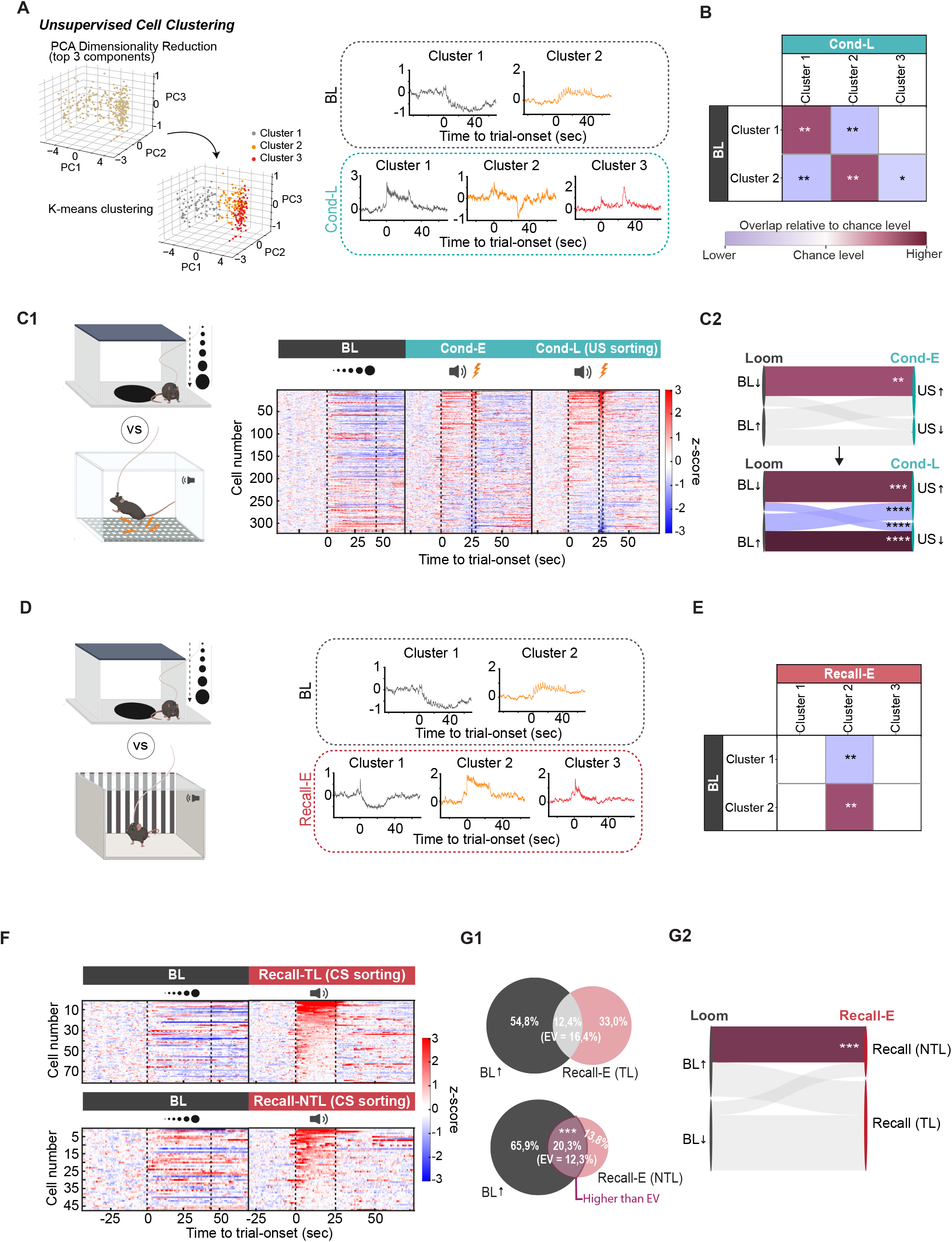
A shared non-time-locked LA ensemble links innate and learned aversive responses. (A) Unsupervised classification of loom- and conditioning-activated neuron dynamics. (Left) The neural activity was reduced with PCA and neurons were grouped with k-means clustering (Loom, k = 3, cluster 1=163 cells, cluster 2=144 cells; Cond-L, k = 7, cluster 1=55 cells, cluster 2=93 cells, cluster 3=78 cells; clusters with little cell numbers <10 or with no significant values are not shown); (Right) Cluster-average traces are shown for the looming session (top) and the late conditioning session (bottom). (B) Contingency matrix between Loom and Cond-L clusters; colors indicate direction of overlap (bordeaux, higher than chance; purple, lower than chance). Overlap significance vs. hypergeometric chance expectation; *p < 0.05, **p < 0.01, ***p < 0.001, ****p < 0.0001. (C) Loom-responsive neurons show anti-overlap with US-responsive neurons during conditioning. (C1) Heatmaps of concatenated single-cell z-scored activity to the cues for the Loom, Cond-E and Cond-L sessions. The cells’ activity is sorted by the response to the US on Cond-L. Dotted vertical lines indicate stimuli onset and offset. (C2) Alluvial diagram tracking the relationship between individual cells from loom-defined groups (BL↑, activated; BL↓, suppressed) to the US-defined response category (US↑, US↓), during Cond-E (top) and Cond-L (bottom). As conditioning progresses from early to late, loom-activated (BL↑) cells become preferentially US-suppressed (US↓), and vice versa, indicating an anti-overlap between BL- and US-responsive populations. Overlap significance vs. hypergeometric chance expectation; **p < 0.01, ***p < 0.001, ****p < 0.0001. (D) PCA and k-mean clustering applied to loom- and recall-activated cells (Loom, k = 3, cluster 1=163 cells, cluster 2=151 cells; Recall, k = 5, cluster 1=148 cells, cluster 2=44 cells, cluster 3=116 cells; clusters with no significant values are not shown). (E) Contingency matrix between Loom and Recall clusters; overlap significance vs. hypergeometric chance expectation; **p < 0.01; asterisks denote greater-than-chance overlap (bordeaux) and significantly less overlap than expected (purple). (F) Heatmaps of single-cell z-scored activity for loom and recall sessions, sorted by the response to the conditioned tone (CS) of the TL neurons (top), and NTL neurons (bottom). Dotted vertical lines indicate stimuli onset and offset. (G1) Venn diagrams showing overlap between loom-activated (BL↑) cells and CS-responsive cells during recall, split into time-locked (TL, top) and non-time-locked (NTL, bottom) subsets. Loom-activated neurons overlapped significantly with the NTL subset (20.3% shared vs. EV = 12.3%; ***p < 0.001), whereas overlap with the TL subset fell below chance at a trend level (12.4% shared vs. EV = 16.4%; p = 0.06). (G2) Alluvial diagram tracking the relationship between individual cells from BL to TL and NTL Recall subgroups. Loom-activated (BL↑) cells overlap preferentially with the NTL population (***p < 0.001).

This observation strongly suggests an anti-overlap between BL-responsive and US responsive neurons. To test this directly, first we sorted all the neurons according to the magnitude of their response to the US. Consistent with the data from the unsupervised clustering, we observed an opposite response direction to the two aversive stimuli: neurons activated by the US were suppressed by the BL, whereas neurons inhibited by the US were activated by the BL (Fig. 4C). Notably, this opposite directionality became more evident with each successive conditioning trial: the neurons activated at the later trials of conditioning were more likely to be suppressed by the BL, and the neurons suppressed during the final trials of the Conditioning were more likely to be activated by the BL (Fig. 4C1,C2; Supplementary Fig. 2A-B).

Animals’ behavioral response to an aversive looming stimulus closely resembles the response to a fear memory recall. In both instances, freezing is the prominent defensive response, and successive stimulus presentations reduce it (habituation and extinction, respectively). This led us to investigate the relation between the neurons encoding the conditioned stimulus versus those encoding the looming stimulus (Fig. 4D). Here, neurons were classified using the same unsupervised method described above, grouping them by their projection into a three-dimensional PC space without prior assumptions about response type.

The most salient feature was the significant overlap between neurons in the BL-activated cluster and those in the cluster showing elevated activity during and after the CS period. Importantly, this correspondence was absent when comparing Loom and Tone sessions, suggesting that the shared population reflects fear memory encoding rather than a general response to auditory stimuli (Supplementary Fig. 2C).

Critically, this cluster-level overlap did not hold at the single-cell level: CS-activated neurons defined by individual response criteria (Fig. 1D) showed no significant overlap with BL-activated neurons (Supplementary Fig. 2D-E). This raises the possibility that within the CS-activated population, a distinct group of neurons is responsible for the observed overlap. Indeed, as we showed earlier, there is a subpopulation of CS-activated neurons — non-time-locked — that undergoes a significant increase in response following conditioning (see Fig. 2F). This subpopulation is a natural candidate for the neurons driving the cluster-level overlap between the Recall-activated and BL-activated clusters. We therefore examined whether a neuron’s response to the BL is related to its pulse-selectivity to the CS (Fig. 4F-G).

We observed that TL neurons activated during the CS were less likely to be activated by the BL stimulus, a trend that approached significance (p = 0.06); on the other hand, NTL neurons were significantly more likely to be BL-activated (Fig. 4F-G). Specifically, about 60% of the NTL neurons were also activated during the BL stimulus (Fig. 4G1). Finally, we asked whether this overlap was a pre-existing property of these neurons or a consequence of conditioning.

Whereas BL-activated neurons overlapped strongly with the NTL population at recall (p = 0.001), no such correspondence was present during the tone on the habituation day, before any tone–shock pairing (p > 0.99; Supplementary Fig. 2F). The shared BL–NTL representation was thus not intrinsic to these neurons but emerged only once the tone had acquired aversive significance, consistent with the idea that this population comes to encode aversive events irrespective of their sensory modality.

### An SST disinhibitory gate controls both innate and learned defensive responses in the LA

The role of inhibitory neurons, particularly somatostatin (SST) and vasoactive intestinal peptide (VIP)-expressing interneurons, has been widely studied in fear learning^40–45^. The dominant model proposes that the activity of excitatory neurons is gated through inhibition by SST neurons (Fig. 5A)^41,43^. As such, input to the excitatory neurons from a neutral tone cannot effectively activate the defensive circuit. During conditioning, the US activates SST-targeting VIP neurons which consequently results in synaptic plasticity. Only after the plasticity, the tone— not neutral anymore— can activate the defensive circuit. We reasoned that the innately aversive looming stimulus bypasses the need for plasticity in part by directly activating VIP neurons. This in turn inhibits SST neurons, ultimately activating the defensive circuit within the LA.

**Figure 5.**
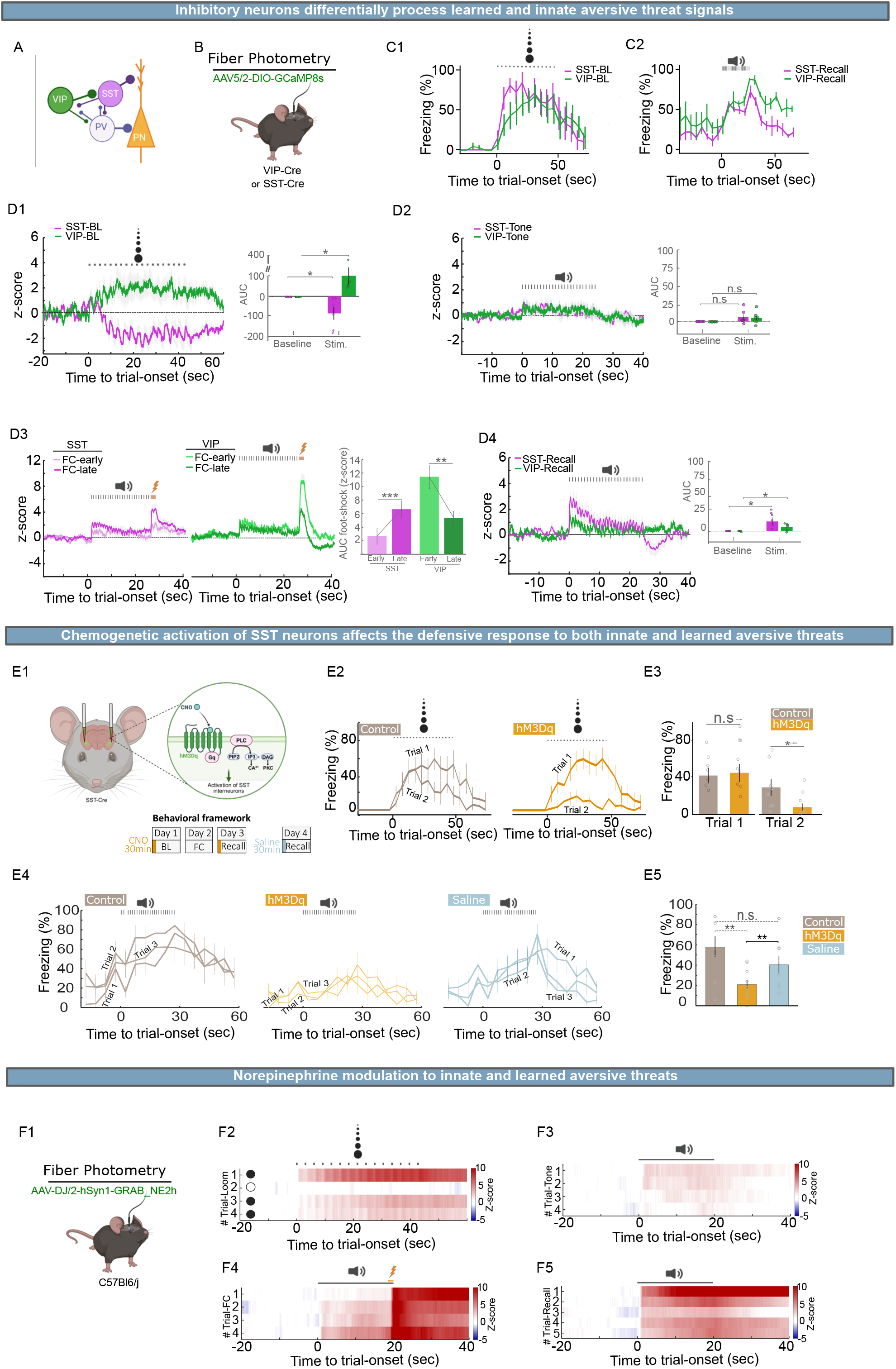
A VIP–SST disinhibitory circuit gates innate and learned defensive responses in the LA. (A) Working model of the LA microcircuit, in which PN activity is gated by a VIP→SST disinhibitory motif: VIP interneurons suppress SST interneurons, relieving their inhibition of PNs (and acting alongside direct PV-mediated inhibition of PNs), thereby permitting PN activation during salient/aversive stimuli. (B) Fiber-photometry strategy: Cre-dependent jGCaMP8s (AAV2/5) expressed in VIP-Cre or SST-Cre mice, with an optic fiber implanted over the LA. (C) Freezing behavior in SST- (n=6 mice) and VIP-Cre (n=8 mice) cohorts. (C1) Freezing to the black-looming (BL) stimulus. (C2) Freezing to the CS during recall. The dotted lines represent stimulus periods. (D) Mean z-scored SST (n=6 mice) and VIP (n=8 mice) bulk activity, with corresponding area-under-the-curve (AUC) quantification (baseline vs. stimulus), for: (D1) the BL stimulus (VIP activated, SST suppressed; t-test, SST, p = 0.043; VIP, p = 0.044); (D2) the neutral tone (n.s.); (D3) the US across early versus late conditioning trials (paired t-test, SST, early vs. late, p < 0.001; VIP, early vs. late, p < 0.01); and (D4) the CS during recall (paired t-test, SST, p < 0.05; VIP, p < 0.05). (E) Chemogenetic activation of SST neurons and its effect on innate and learned defensive responses. (E1) Strategy and experimental design. Excitatory hM3Dq DREADD was expressed in SST neurons of SST-Cre mice; a separate SST-Cre cohort expressing GFP served as control. hM3Dq mice received CNO 30 min before the looming session and 30 min before recall; the following day, the same mice received saline 30 min before a second recall session (within-subject CNO vs. saline). Control (GFP) mice received CNO 30 min before looming and recall (n_hM3Dq = 11 mice, n_control = 8 mice). (E2, E3) Freezing to the BL stimulus across trials 1–2, comparing control (GFP + CNO) and hM3Dq + CNO mice (between-subjects). hM3Dq + CNO mice show reduced freezing on trial 2 (unpaired t-test per trial: Trial 1, n.s.; Trial 2, *p < 0.05). (E4, E5) CS-evoked freezing at recall for control (GFP + CNO), hM3Dq + CNO, and hM3Dq + saline mice. The hM3Dq + CNO and hM3Dq + saline conditions are the same animals on consecutive days (within-subject, paired), whereas the control cohort is separate (between-subjects, unpaired): control vs. hM3Dq + CNO, unpaired, p < 0.01; hM3Dq + CNO vs. hM3Dq + saline, paired, p < 0.01; control vs. hM3Dq + saline, unpaired, n.s. Error bars represent s.e.m. (F) Norepinephrine (NE) dynamics in the LA during innate and learned aversive stimuli. (F1) Strategy for imaging NE release with fiber-photometry using the GRAB-NE2h sensor. (F2) NE is released in response to black-looming but not white looming, and this response declines across trials as animals are exposed to subsequent BL (n=17 mice). (F3) NE release to the neutral tone (habituation) is minimal (n=15 mice). (F4) NE is released in response to the US during conditioning; and across conditioning trials, the tone itself progressively comes to elicit NE release (n=15 mice). (F5) During recall, the CS now elicits NE release, which subsequently tends to decline across trials as the CS is no longer reinforced (n=14 mice).

To test this, we recorded bulk calcium activity in VIP- and SST-Cre mice expressing Cre-dependent GCaMP8m in the LA. GCaMP signal was collected through a fiber optic implanted in the LA (Fig. 5B). Animals underwent the same behavioral paradigm used for microendoscopic calcium imaging, recording the activity during Habituation, Conditioning, Recall, and BL exposure (Fig. 5C).

The BL stimulus had a significant and opposing impact on the activity of these two inhibitory neurons: it increased the activity of disinhibitory VIP neurons while suppressing the activity of inhibitory SST neurons (Fig. 5D1). This change outlasted the BL offset for many seconds. The neutral tone, on the other hand, had only a marginal effect on either interneuron population (Fig. 5D2). In contrast to the BL stimulus, neither the disinhibitory VIP neurons nor the inhibitory SST neurons changed their activity significantly from baseline during the tone, and the modest fluctuations that were present did not outlast the stimulus. Together, these data indicate that the innately aversive BL and the neutral tone are read out differently by LA interneurons: the BL, but not the neutral tone, drives the VIP activation and SST suppression, opening the gate to the defensive circuit (Fig. 5D1,D2).

We next followed how these interneurons behaved across the course of conditioning (Fig. 5D3). At training onset, the US evoked a sharp transient response in both the inhibitory SST and the disinhibitory VIP neurons, with the VIP response markedly the larger of the two. As conditioning progressed, however, the two populations moved in opposite directions: the US-evoked response of SST neurons grew from the early to the late conditioning trials, whereas that of VIP neurons declined over the same trials, so that their relative magnitudes had effectively reversed by the end of training (Fig. 5D3).

At recall, the tone — ineffective before training — now engaged the inhibitory circuit, inducing a transient activation of SST and VIP neurons at tone onset (Fig. 5D4). Notably, this initial SST activation was biphasic: at the trial offset, it gave way to a suppression of SST activity below baseline that outlasted the tone (Fig. 5D4). These recordings show that SST suppression accompanies the defensive response to both innate and learned threats, but they do not establish that it is required for either.

To test this causally, we expressed the excitatory DREADD hM3Dq^46^ in SST neurons of SST-Cre mice and chemogenetically forced SST activity by injecting CNO 30 minutes before the looming session and, two days later, before the recall session; a separate recall session preceded by saline injection served as a with-in-subject control (Fig. 5E1, Supplementary Fig. 3). Another group of animals expressing GFP instead of hM3Dq served as a control for the effects of CNO and the DREADD manipulation itself, providing a baseline for both the BL and Recall sessions.

We argued that if SST suppression is necessary to open the defensive circuit, then forcing SST activity should attenuate defensive responding. This is what we observed. Although hM3Dq mice showed a normal response to the first BL presentation, their freezing rapidly collapsed by the second trial, falling well below the gradual habituation seen in control animals (Fig. 5E2,E3).

The same manipulation impaired the learned defensive response with activating SST neurons reduced the CS-evoked freezing at recall relative to the control mice (Fig. 5E4,E5). Importantly, the freezing response to the CS was restored in a subsequent session in which hM3Dq-expressing animals received saline, indicating that the deficit was CNO-dependent rather than a consequence of hM3Dq expression or repeated testing. Thus, elevated SST activity is sufficient to attenuate both innate and learned defensive responses in the LA.

We also recorded PV neurons under the same conditions (Supplementary Fig. 4). Unlike SST and VIP neurons, PV neurons were activated in the same direction by every stimulus we tested — the BL, the US, and the tone at recall — and, unlike either population, they were also activated by the neutral tone (Supplementary Fig. 4C1–C4). Their US response, moreover, did not change between early and late conditioning, showing none of the reorganization seen in SST and VIP neurons (Supplementary Fig. 4C3). Thus PV neurons track aversive and neutral stimuli alike in a stable, non-selective manner, rather than participating in the stimulus- and learning-specific gating that distinguishes the SST and VIP populations.

### Norepinephrine release tracks innate and learned threats in the LA

Previously, we showed that systemic inactivation of the β-adrenergic receptor blunts the animals’ defensive response to BL stimuli and reduces the LA neuronal response to the stimuli^26^. This suggests that the innately aversive visual stimulus triggers the release of norepinephrine (NE) in the LA. To directly monitor NE release in the LA, we used the genetically encoded sensor GRAB_NE2h^47^ (Fig. 5F1) during Habituation, Conditioning, Recall, and BL presentation.

The innately aversive BL stimulus caused a significant release of NE in the LA; in contrast, the neutral WL had no effect. Across successive trials with the BL stimuli, the level of NE release dropped as the mice habituated (Fig. 5F2). Compared to the BL, the neutral tone triggered only a modest release of NE, primarily during early trials when the tone was novel (Fig. 5F3). However, during conditioning, the US triggered a massive release of NE (Fig. 5F4), and notably, we observed an increasing release of NE to the tone as conditioning progressed. In the recall session, once the tone became a predictor of a US, it caused a considerable release of NE (Fig. 5F5).

Together, these data show that NE release in the LA follows the same pattern as SST suppression: present for the innately aversive BL, absent for neutral stimuli, and appears for the tone over conditioning. This raises the possibility that neuromodulatory input contributes to the interneuron dynamics described above, a link that remains to be tested directly.

## Discussion

The main objective of this work was to compare side-by-side the circuits processing innate and learned threats in the LA. This was motivated by the fact that both types of stimuli activate the same region and induce similar defensive responses. It would, therefore, be evolutionarily parsimonious if the two circuits shared common elements^26^. Identifying such shared elements could, in turn, provide new insight into the cellular mechanisms of learning.

The three core elements of the circuit processing learned aversive responses in the LA— excitatory neurons, inhibitory neurons, and neuromodulators— are well studied. Our findings largely corroborate previous works, but with some notable refinements. It has been reported that following conditioning, around a third of CS-activated neurons increase their response to the CS, while the response of another third remains stable^35^.

Here, we show that the neurons that increase their response amplitude following conditioning are distinct from stable neurons at least in two features. First, prior to conditioning, the neurons that later increased their CS-response had a smaller response amplitude to the tone than the stable neurons. This may reflect a weaker synaptic connection to their upstream, tone-selective neurons before training. Second, the two groups differed in their tone-response dynamics: the neurons with sustained activity throughout the pulsated tone period (NTL) were significantly more likely to increase their response to the CS during and following conditioning.

This group of neurons appears to be a link between the circuits processing innately aversive looming stimulus and learned cued conditioning. Specifically, the neurons that were activated by the looming stimulus were significantly more likely to belong to the NTL population. Conversely, the tone-pulse–locked (TL) neurons showed the opposite tendency, being suppressed rather than activated by the looming stimulus—an anti-overlap that approached significance (p = 0.06).

This suggests that the neurons recruited for aversive learning during conditioning may not be selected randomly. Although at this stage we do not know the origin of TL and NTL neurons, we suspect that they are differentially driven by the two major sources of auditory inputs to LA. Specifically, TL neurons may be driven predominantly by upstream cortical inputs, while NTL neurons may receive input from thalamic regions^48^. This attribution is consistent with previous work: inactivating thalamic inputs to the LA impairs both the innate and learned defensive responses, whereas inactivation of cortical inputs has no behavioral effect. This parallels the NTL population, which is engaged by both the innately aversive loom and cued conditioning^20,26^.

The second link between the circuits processing innate and learned defensive responses is inhibitory neurons. The canonical model proposes that synaptic plasticity mediating the learning is gated through the disinhibitory VIP-SST microcircuit ^41,43^. Our data indicate that, unlike a neutral tone, an innately aversive looming stimulus engages this disinhibitory gate directly—activating the VIP neurons and thereby suppressing SST neurons—without requiring the learning-related plasticity that a neutral tone depends on. Supporting this, artificial activation of SST neurons in the LA not only impairs learned defensive response, but also induces rapid habituation to the innately aversive looming stimulus. On the other hand, activation of SST neurons in the superior colliculus— a region essential for the innate but not the learned defensive response— has no effect on the animals’ defensive response^49^.

This dissociation indicates that the looming response sensitivity to SST-mediated inhibition is a specific property of the learning-capable LA circuit rather than a general feature of innate threat processing. These findings suggest that both the CS and aversive looming stimulus tap into the same microcircuitry interneurons in the LA, with the innate stimulus having privileged access, while the learned cue relies on synaptic plasticity.

Before proceeding further, two puzzling observations must be addressed. First, while both looming stimulus and US are aversive, we see an anti-overlap between the neuronal populations activated by the two stimuli. Moreover, as conditioning progresses, this anti-overlap becomes more pronounced. Although at this stage we do not grasp the biological significance of this phenomenon, we speculate mechanistically that the anti-overlap may be mediated by SST neurons because the black looming stimulus suppresses SST neurons, whereas the US activates them, especially as conditioning progresses.

Second, two observations seem to be in contradiction: on one hand, during the recall session, CS activates SST neurons. On the other hand, artificial activation of SST neurons during the recall session impairs the freezing response. These two findings can be reconciled by the reports that there are two populations of SST neurons in the LA with opposing responses to the CS^44,45^. Although our bulk calcium imaging cannot resolve these two populations during the CS, we observe an immediate suppression of SST activity following the CS offset, perhaps representing the neurons whose suppression permitting the expression of the defensive response, was masked during the tone by CS-activated SST neurons.

The CS-suppressed SST neurons may belong to the class that was suppressed by the looming stimuli, a class whose activation reduces the defensive response to both innate and learned aversive stimuli (Fig. 5E). This suggests that suppression of these neurons during aversive experience disinhibits the defensive circuit to promote the freezing response. The CS-activated SST neurons, by contrast, may sharpen the memory trace: by laterally inhibiting neighboring principal neurons, they could restrict the number of cells recruited into the engram, constraining its size while enhancing its stimulus specificity—a role analogous to that reported for SST neurons in the hippocampus and PV neurons in the LA^50,51^.

The third and final link that we explored was the neuromodulator, norepinephrine (NE). Both the innately aversive stimulus and the learned aversive stimulus triggered significant release of NE which outlasted the stimulus. The notable difference was that, unlike for the innate aversion, the tone acquired this capacity only through learning, as it had little effect before conditioning. This aligns with other works showing that innate defensive responses to predator odors or looming stimuli in mice require the release of NE^26,52–55^. Similarly, both the formation and stability of an aversive associative memory in the LA require the activation of β-adrenergic receptors^10,56^. In both forms of aversive experiences, the neuromodulator may act through similar cellular mechanisms: enhancing the excitability of the pyramidal neurons either directly by downregulating potassium channels in these neurons^57^; or indirectly by reducing excitability of inhibitory neurons^58^.

Taken together, these results suggest that the circuit processing learned aversive experience recruits a circuit in the LA that evolved to protect animals from their natural predators. This could explain why animals learn to recognize new threats so quickly^59^. Such interactions between circuits underlying innate and learned behavior are unlikely to be unique to the LA^60^. Co-opting an existing circuit would conserve space and energy that would be needed to create a new circuit de novo.

## Methods

### Animals

All animal procedures were performed in accordance with the institutional guidelines at the Department of Molecular Biology and Genetics, and the Department of Biomedicine, at Aarhus University, and approved by the Danish Animal Experiment Inspectorate under the permit numbers 2020-15-0201-00421 and 2023-15-0201-01431.

C57BL/6JRJ wildtype male mice (Janvier, France) were used for Miniscope experiments, and for fiber photometry experiments involving norepinephrine sensors. VIP-Cre, SST-Cre and PV-Cre mice (males and females, local breeding) were used for Cre-dependent expression of viral vectors for fiber photometry calcium imaging and for chemogenetic manipulation. All animals were aged 2-3 months by the time of the virus injection. The animals were group-housed (3-4 per cage) with enriched conditions in a 12h light/dark cycle, with food and water *ad libitum*. The animals implanted with a GRIN lens for Miniscope calcium imaging were single-housed with enriched conditions, from the implantation day until the end of the experiments. All behavioral experiments were conducted during the light cycle.

### Deep-brain calcium imaging

#### Stereotaxic surgeries and virus expression

Mice were anesthetized using a cocktail of Fentanyl, Midazolam and Medetomidine (FMM, injected intraperitoneally (i.p.)). Before the surgery, the mice were injected subcutaneously with Buprenorphine 0.3 mg/ mL (Temgesic, 0.1 mg/kg). The use of stereoscope lights was kept minimal to prevent damaging the eyes and ophthalmic ointment (Viscotears, Bausch and Lomb) was applied regularly to prevent eye drying. The body temperature was maintained at 37°C during the surgery.

Once anesthetized, mice were fixed to a stereotaxic apparatus (KOPF Instrument, model 963). Standard surgery procedures were used to expose the skull and drill a craniotomy above the target site.

AAV1/2-mCaMKIIα-jGCaMP8m (900nL, University of Zurich facility, VVF; 6.8 x 10E12 vg/ml diluted 1:4 in sterile PBS) was unilaterally injected using a nanoliter injector Nanoject III (Drummond, 2nL per second) and pulled glass pipettes at the following coordinates: antero-posterior from bregma (AP) -1.7 ; medio-lateral from bregma (ML) ; -3.45/-3.55 ; dorso-ventral from the bone, from -5.1 to -4.1 with 150nL of virus injected every 0.2 mm. Following the virus injection, the craniotomy was sealed with fast-drying silicone (Kwik-Sil, World Precision Instruments), and the skin was closed with staples.

Between three to five weeks after virus injection, a GRIN lens (ProView Integrated lens 0.5 × 6.1 mm, Inscopix) was implanted into the LA. In brief, a sterile needle (20G) was used to make an incision above the imaging site. The GRIN lens was subsequently lowered into the brain using a stereotaxic motorized manipulator (Scientifica IVM, coordinates: AP: –1.7 mm; ML: –3.55 mm; DV from bone: 4.9 mm). The implantation was done under visual inspection by connecting the integrated lens on the miniscope secured by a holder. Once in place, the lens was fixed to the skull using light-curable adhesive (OptiBond, Kerr), followed by light-curable cement (VertiseFlow, Kerr). After the surgery, the animals received an i.p. injection of antidote (a mixture of Naloxone, Atipamezol and Flumazenil) and a postoperative treatment with Temgesic in drinking water for 3 days. Animals were left to recover for 3 weeks after GRIN lens implantation before checking for GCaMP expression. The behavioral experiments were performed from 4 to 7 weeks post-implantation, or until the tissue was healed from the surgery.

#### Data Acquisition

Starting 4 weeks following GRIN lens implantation, the miniscope was mounted to check for GCaMP expression as well as for healing and stability of the tissue, and the animals were acclimated to carry the miniscope in an open-field. The miniscope was removed after each imaging-session. Imaging data was acquired using Inscopix Data Acquisition Software at a frame rate of 20 Hz with an LED power of 0.1-0.5 (475 nm), analog gain of 0.3–1 and a field of view set where most cells were sharp (usually 350-700 units). The same imaging parameters were kept across repeated behavioral sessions for each individual mouse. To reduce bleaching during the behavioral experiments, the recordings started approximately 3 minutes prior to the presentation of the first stimulus.

#### Image processing and analysis

#### Pre-processing

First, raw images were cropped to the circular field of view delimited by the visible GRIN lens track, excluding peripheral optical distortion and spurious objects (e.g., debris) at the edge of the field of view. Then, to extract calcium activity from the imaging sessions, the videos were spatially and temporally downsampled by a factor of 2. A spatial bandpass filter was then applied to remove background fluorescence and reduce noise. Motion artifacts were corrected using a frame-to-frame image registration approach with a selected reference frame. These preprocessing steps were performed using Inscopix Data Processing Software (IDPS).

Because recordings from the amygdala contained substantial motion artifacts and blurring, automated segmentation methods such as CNMF-e were not reliable. Therefore, regions of interest (ROIs) were manually defined on motion-corrected and ΔF/F videos, as well as on maximum projection images, using a custom Python-based tool, **PickCell** (Supplementary Fig. 5A1). ΔF/F was calculated for each pixel as ΔF/F = (F(*t*) − F_0_) / F_0_, where F(t) is the pixel intensity at time *t* and F_0_ is the mean pixel intensity across the session. Calcium activity was then extracted by averaging the ΔF/F values within each ROI over time.

To identify the same neurons across recording sessions, ROI maps were aligned using rotation and translation to maximize spatial correspondence. This alignment procedure was also performed using **PickCell** (Supplementary Fig. 5A2).

#### Activity normalization

The activity of each cell during a trial was normalized to its own baseline using a z-score, calculated as *z*_*i*_*(t) = (x*_*i*_*(t)* − *μ*_*i*_*) / σ*_*i*_, where *x*_*i*_*(t)* is the raw activity (ΔF/F) of cell *i* at time *t*, and *μ*_*i*_ and *σ*_*i*_ are the mean and standard deviation of that cell’s activity during the 60 seconds preceding trial onset, respectively.

#### Cells classification

To identify stimulus-responsive cells, a selectivity analysis was performed by comparing each cell’s activity during the stimulus period with its baseline activity on a trial-by-trial basis. First, the normalized activity was averaged into time bins (0.2 s bins for US and 1 s bins for other stimuli). A Wilcoxon signed-rank test was then used to assess significant deviations from baseline (p < 0.05).

Cells showing a significant increase in activity were classified as Up (excited), cells showing a significant decrease were classified as Down (inhibited), and all remaining cells were classified as None (non-responsive). To identify responses that were consistent across trials, bin-averaged activity values from the stimulus and baseline periods were pooled across trials before applying the statistical test.

To identify neurons potentiated to a stimulus from one session to another, we compared their z-scored responses to the stimulus between the two sessions. Potentiation was assessed per neuron by comparing the area under the curve (AUC) of the CS-evoked response between the two sessions. Neurons whose AUC in the second (Recall) session was at least twice their AUC in the first (Tone) session were classified as potentiated; all remaining neurons were classified as non-potentiated. .

#### Functional similarity analysis

To measure functional similarity between neurons, we calculated pairwise cosine similarity scores for all pairs of neurons based on their trial-averaged z-scored activity across sessions as s_ij_ = (z_i_ · z_j_) / (|z_i_| × |z_j_|), and z_i_ and z_j_ are the trial-averaged z-scored activity vectors of neurons i and j, respectively.

#### Principal Component Analysis (PCA)

To reduce data dimensionality and visualize neural population dynamics across sessions, principal component analysis (PCA) was applied to trial-averaged population activity from selected trials across one or multiple sessions. For each session, the trial-averaged activity of tracked neurons was arranged into a matrix, with time points as rows and neurons as columns.

These matrices were concatenated along the time dimension to create a pooled activity matrix, which was used for PCA to identify a common low-dimensional subspace across sessions. The first three principal components were then used to project the trial-averaged activity from each session, generating low-dimensional neural trajectories over time. (Yonehara et al., 2023)

#### Pulse-selectivity analysis

To identify cells responsive to tone pulses, we developed a pulse-selectivity analysis (Fig. 2E1). Tone pulses were 200 ms in duration, and the response window was defined as 0–0.4 s after pulse onset to account for the slower kinetics of calcium signals. Cells were classified as time-locked (TL) if the slope of their activity during the response window differed significantly from the slope during the inter-pulse interval (0.4–1 s after pulse onset; p < 0.05).

#### Overlap significance

To test whether the overlap between two cell populations was greater or less than expected by chance (chance level), the expected overlap value (EV) was calculated as EV = (M_1_ × M_2_) / N, where M_1_ and M_2_ are the numbers of cells in populations 1 and 2, respectively, and N is the total number of cells.

Statistical significance was assessed using a two-sided Fisher’s exact test based on the hypergeometric distribution. The p-value was calculated by summing the probabilities of all possible overlap values with probabilities equal to or lower than the probability of the observed overlap.

A p-value below 0.05 was considered significant. Significance levels are indicated as *p < 0.05, **p < 0.01, ***p < 0.001, ****p < 0.0001.

#### Contingency matrix

To evaluate the significance of overlap between all response populations for two stimuli, a contingency matrix was constructed. The rows and columns represent the response categories for each stimulus, and each entry shows the statistical significance of the overlap between the corresponding response populations.

#### Unsupervised cell clustering

Similar to the selectivity analysis, we also applied an unsupervised approach to identify groups of cells with similar response profiles. For each cell, activity was averaged across the corresponding trials, yielding an N × T matrix, where N is the number of cells and T the number of time points. We reduced dimensionality by principal component analysis (PCA), projecting each cell onto the first three principal components (N × 3) to capture the largest sources of variance. k-means clustering was then performed in this three-dimensional space. The number of clusters was set a priori for each session (k = 3 for Loom, k = 6 for late conditioning [Cond-L], and k = 5 for Recall), chosen by visual inspection of the cluster-average dynamics to capture the distinct response profiles present in each session without over-fragmenting them. The results were robust to the exact choice of k.

### Fiber photometry of inhibitory neurons and norepinephrine release

#### Stereotaxic surgeries and virus expression

Transgenic animals (VIP-cre, SST-cre, and PV-cre) or C57BL/6JRJ wild-type mice were anesthetized either with FMM as described above, or with Isoflurane (IsoFlo vet, Zoetis, 5% for induction, 2% for maintenance). The mice were then fixed onto a digital stereotaxic apparatus (KOPF Instrument, model 940). Standard surgery procedures were used to expose the skull and drill the craniotomy above the target site.

AAV5/2-hSyn-DIO-jGCaMP8s (800nL, 5.8 x 10E12 vg/ ml, University of Zurich facility, VVF), or AAV-DJ/2hSyn1-GRAB_NE2h-WPRE-hGHp(A) (800 nL, custom-made by VVF, gifted by Tomonori Takeuchi lab, Aarhus University, Denmark), was unilaterally injected using a nanoliter injector Nanoject III (Drummond, 2nL per second) or a Picospritzer III microinjection system (Parker Hannifin Corporation) and pulled glass pipettes at the following coordinates: antero-posterior from bregma (AP) -1.7 ; medio-lateral from bregma (ML) ; -3.45/-3.55 ; dorso-ventral from the bone, from -4.9 to -4.1 with 150nL of virus injected every 0.2 mm.

Following virus injection, an optic fiber (300 µm Ø, NA 0.37, RWD) was implanted into the LA at the following coordinates: AP: –1.7 mm; ML: –3.55 mm; DV from bone: -4.3 mm and fixed to the skull using light-curable adhesive (OptiBond, Kerr), followed by light-curable cement (VertiseFlow, Kerr). After 5 to 8 weeks post-surgery, the animals were handled for two to three days before the beginning of the experiments.

#### Data Acquisition

All the recordings were performed with a Doric fiber photometry system composed of an LED-driver, a fiber photometry console, and a Doric mini-cube with 460-490 nm for GCaMP excitation and 415 nm for isosbestic excitation, and a low-autofluorescence patch cord (200 nm or 300 nm, 0.37 NA) was used. The latter was bleached for at least 5 hours before each experiment via a 473 nm laser at 15-17 mW. The GCaMP signal was amplified with a Doric amplifier with a 10x gain and recorded with Doric Neuroscience Studio software (Version 5.4.1.23) at 11 kHz. Light power at the patchcord tip was set between 60 to 65 µW for 470 nm excitation, and between 30 to 35 µW for 415 nm excitation.

For synchronization with the looming stimulus presentation, a National Instrument board (NI USB 6003) was used to timestamp the looming stimulus presentation over the calcium signal. For synchronization with the tone and shock presentations, an input-output box connected to the aversive conditioning system was used.

#### Data processing

Fiber photometry data were processed using a custom MATLAB script to extract calcium activity from neural recordings. First, the raw calcium (GCaMP) and reference (isosbestic) signals were smoothed using a median filter (`medfilt1`) with a 0.1-second window. To remove initial recording artifacts, the first 5 seconds of data were discarded. The signals were then downsampled by a factor of 10 using the `decimate` function, reducing the sampling rate from 12,048 Hz to 1,204.8 Hz.

To correct for photobleaching, when present, the calcium signal was detrended using a biexponential fit (`fit(t,x`,’`exp2`’)). To account for motion-related artifacts, the isosbestic signal was scaled to match the amplitude of the calcium signal using a linear fit (`polyfit`). If the fitted line had a positive slope, calcium activity was calculated as 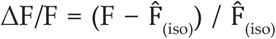, where F is the calcium-dependent fluorescence signal (GCaMP) and 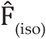 is the fitted and scaled isosbestic reference signal. If the fitted slope was not positive, the signal was baseline-corrected by subtracting its mean value.

Finally, to enable comparisons across animals, the activity was normalized on a trial-by-trial basis using a z-score, calculated from the mean and standard deviation of the pre-trial baseline period.

All the traces with a sudden change in the isosbestic signal were discarded in the final analysis.

For Tone, Conditioning and Recall, we considered the first 10 seconds after CS onset in the analysis (Fig. 5D2D4).

### Behavioral paradigm

#### Looming stimulus

The apparatus consisted of an opaque open-top arena (37×40×19.5 cm) with a monitor (16 inches) placed on the top. No shelter was used^30,61^. The arena was placed in a dark room. The animals were placed in the center of the arena and were left to explore freely for approximately 10 minutes prior to the first exposure to the looming stimulus. The looming stimulus consisted of 15 repetitions, from 2° to 20° of visual angle^27^ of expanding black disk over a gray background. Each loom widened in 800 ms and remained at the same size for 250 ms, followed by 2 seconds of pause before the next loom appeared. An interval of minimum 3 minutes separated consecutive trials. A white looming stimulus (same properties), was then delivered to serve as a control for the lack of behavioral reaction to a non-aversive visual stimulus. The behavioral responses were recorded with an infrared top camera (Phihong POE21U-1AF). The animals’ time spent in freezing was quantified manually using a self-made software (Supplementary Figure 5B). Freezing was defined as a complete lack of movements, except for the respiratory movements, that lasted for at least 1s.

#### Aversive conditioning and retrieval

Two different contexts were used for the associative learning paradigm. Context A (pure tone and retrieval) consisted of a patterned-wall open-top box (24×20×30 cm) with a smooth floor placed into a soundproof cubicle (55×60×57 cm) (Ugo Basile, Italy) with ambient odor (isoamyl-acetate). Context B (threat conditioning) consisted of a clear open-top box (24×20×30 cm) with an electrical grid floor for footshock delivery placed into the soundproof cubicle.

For both contexts, a speaker delivered a custom-made auditory stimulus, triggered by ANY-maze software (Stoelting, Ireland). The auditory stimulus consisted of an epoch of 25 x 200 ms pips repeated at 1Hz (CS: 5kHz) at 70 dB. For the experiments involving NE neurosensors the auditory stimulus consisted of 20-sec of constant tone (5khz, 70dB)

On day 1, mice were habituated to context A for 3 minutes prior to the delivery of 5 trials of the auditory stimulus, presented with a pseudorandom inter-trial interval (ITI, range of 60-120s).

On day 2, mice were conditioned in context B to associate the CS with a US (2 sec of 0.6 mA footshock) delivered upon the termination of the auditory stimulus, over 5 trials with pseudorandom ITI. After the conditioning, the animals, if group-housed, stayed isolated for 10-15 minutes before returning to their homecage.

On day 3, the retrieval of the memory was assessed in context A. After 3 minutes of free exploration, the mice were exposed to 12 trials of non-reinforced CS.

#### Behavioral scoring

Animal behavior during recall sessions was scored using ANY-Maze by measuring the percentage of time spent freezing in 5-second bins, with a minimum freezing duration of 1 second (as defined in the software). For Looming sessions, mouse behaviors, including freezing, rearing, and tail-rattling, were manually scored using a custom Python-based tool, **BehaviTrack** (Supplementary Fig. 5B). The tool allows users to review videos at a fine temporal resolution (0.1 s), annotate behaviors, and track the animal’s position in the arena. For these analyses, the minimum freezing duration was set to 0.5 seconds.

#### Chemogenetic manipulation of SST inhibitory neurons

SST-Cre mice underwent surgical procedures as described above. AAV5/2-hSyn-DIO-hM3Dq-mCherry (800nL, 4.0 x 10E12 vg/ml) or AAV-5/2-hSyn-DIO-eGFP (800nL, 1.1 x 10E13 vg/ml) was bilaterally injected using a Picospritzer III at the following coordinates: (AP) -1.7; (ML); -3.45/-3.55; (DV from the bone), -4.4 to -4,1. Following the injection, the skin was closed with surgical clips and the animals received an injection of carprofen before recovering on a warm plate. The behavioral experiments were performed from 5 to 8 weeks post-surgery.

On the experiment day, Clozapine N-oxide dihydrochloride (CNO water-soluble, HelloBio, HB6149; 10 mg/kg, 400 µL) or saline (0.9% NaCl, 400/500 µL) were administered by intraperitoneal injection. After the injection, the mice were isolated for 30 min before the beginning of the experimental procedures.

### Histology and immunofluorescence

Following completion of the experiments, the animals were anesthetized with FMM and euthanized by cervical dislocation. The head with implants was fixed in 4% paraformaldehyde for at least 3 days at room-temperature (RT). After fixation, the implant was removed and the brain section containing the LA was sliced in 300 µm coronal sections using a Vibratome (Leica, VT1000 S). The implant location and virus expression were verified under an upright microscope against a mouse brain atlas. Animals with misplaced implants or improper virus expression were excluded from the analysis.

To rule out any virus-induced toxicity in the implanted animals, the brain slices were stained for NeuN. Free-floating slices were permeabilized and blocked with a mixture of 0.5% TritonX, 10% Normal Goat Serum (NGS) and 10% Bovine Serum Albumin (BSA) in PBS, for 90 min at RT. Subsequently, the slices were incubated for 72h at 4°C, with a mixture of anti-NeuN antibody (Merck Millipore, MAB377; 1:500, in mouse), 0.1% Triton X and 1% NGS in PBS. At the end of the incubation, the slices were washed three times in PBS, followed by an incubation in Cyanine 3 (Cy3, Goat anti-mouse, Thermo Fisher Scientific, A10521, 1:500) with 0.3% Triton X, 1% NGS and 5% BSA in PBS for 24 hours at 4°C. Nuclear staining was performed by using 1:1000 of DAPI (Sigma, D9542) for 30 minutes at room temperature. Brain slices were mounted on polylysine glass slides with coverslips using Fluoromount G (Southern Biotech).

Imaging was performed by using a ZEISS Apotome microscope (Axio Imager.M2) linked to ZEN software (ZEISS). Images were taken from the region of interest using a 10x objective (Apochromat 10/0.45 M27). Excitation wavelengths for GCaMP/GFP and Cy3/mCherry visualization were 518 nm and 565 nm respectively.

Some of the panels in Figures 1,3,4,5 and Supplementary figures 4 and 5 were made using BioRender.com

## Authors’ contribution

NMJ, MN and SN conceived ideas, analyzed all the data and wrote the manuscript. NMJ performed the experiments. AP contributed in acquiring and analyzing photometry data. SA extracted and processed the calcium imaging and behavioral data. MN developed methods for data processing and data analysis. JSL contributed to behavioral data analysis and brain tissue processing. AKV provided technical assistance. All the authors revised the manuscript.

## Acknowledgments

We Thank P. Sterling, R. Malinow, D. Kvitsiani, and J. Piriz for their comments on the manuscript. We thank the current and previous members of the Nabavi lab for their suggestions during the progress of this project. This study was supported by the Novo Nordisk Foundation (NNF0087010); PROMEMO – Center for Proteins in Memory, a Center of Excellence funded by the Danish National Research Foundation (DNRF133); and the Lundbeck Foundation (grant no. DANDRITE-R248-2016-2518, and grant no. R400-2022-1177 to NMJ).

## Data availability

The data generated in this study are available from the corresponding author upon request.

---

**Supplementary Figure 1.**
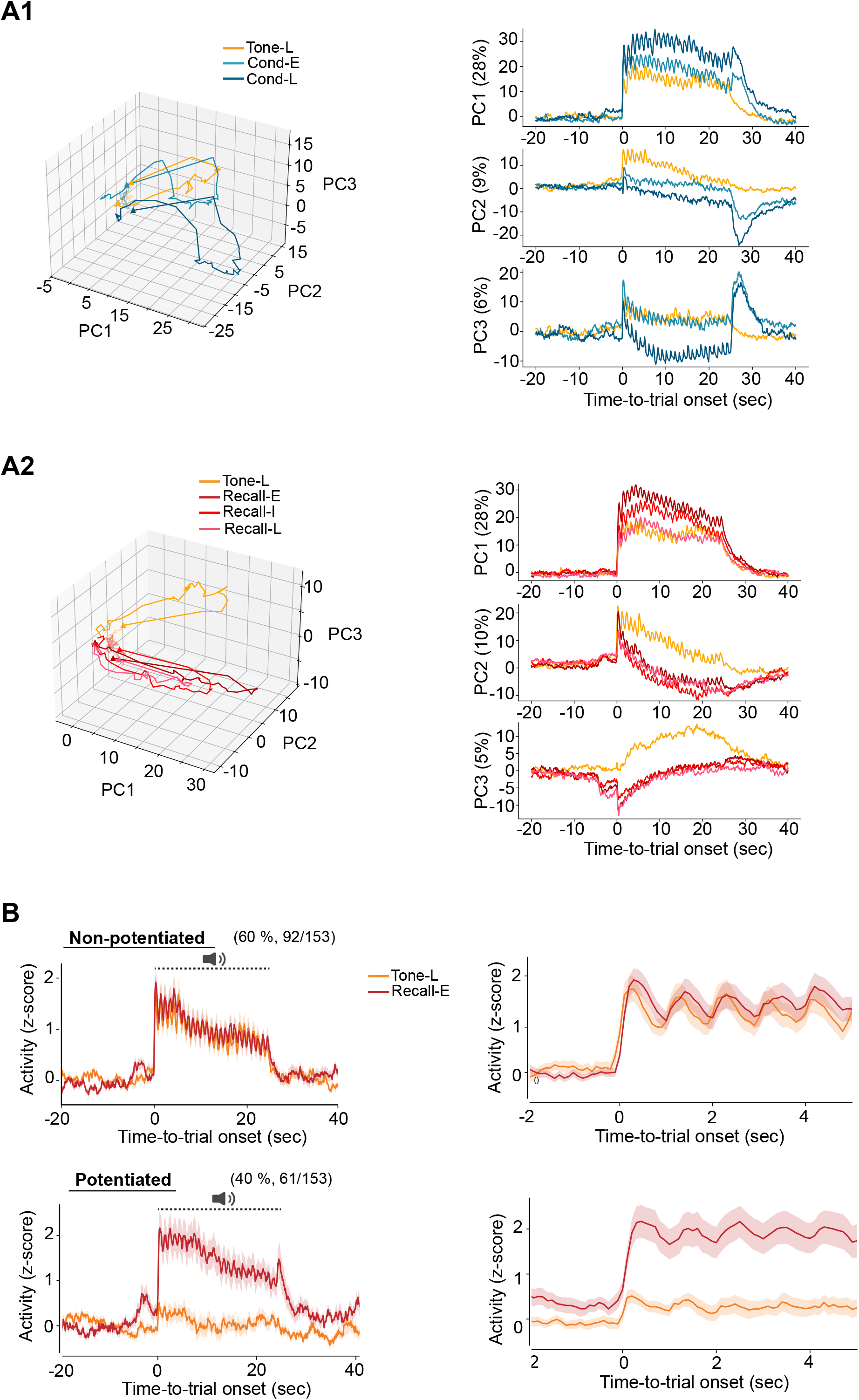
Population dynamics across tone, conditioning, and recall sessions. (A) Principal component analysis (PCA), (A1) Three-dimensional state-space trajectories (PC1–PC3) during tone presentation (Tone-L), early conditioning (Cond-E), and late conditioning (Cond-L), with corresponding individual PC projections (PC1, PC2, PC3) plotted as a function of time-to-trial onset. The percentage of variance explained by each component is shown in parentheses. (A2) As in A1, but comparing tone (Tone-L) to early (Recall-E), intermediate (Recall-I), and late (Recall-L) recall session. (B) Average population activity (z-scored) for Tone-L vs. Recall-E, for neurons classified as potentiated (top) or non-potentiated (bottom) based on the change in their CS-evoked response from the Tone to the Recall session (n = 153 CS-activated neurons). Neurons whose CS-evoked AUC at least doubled from the Tone (Tone-L) to the early recall (Recall-E) session were classified as potentiated (40%, 61/153 neurons); the remainder were classified as non-potentiated (60%, 92/153 neurons) (see Methods). (Left) Full trial time course (−20 to 40 s; dashed-line indicates the stimulus period); (Right) expanded view of the peri-tone-onset window (−2 to 5 s). Shaded traces represent s.e.m.

**Supplementary Figure 2.**
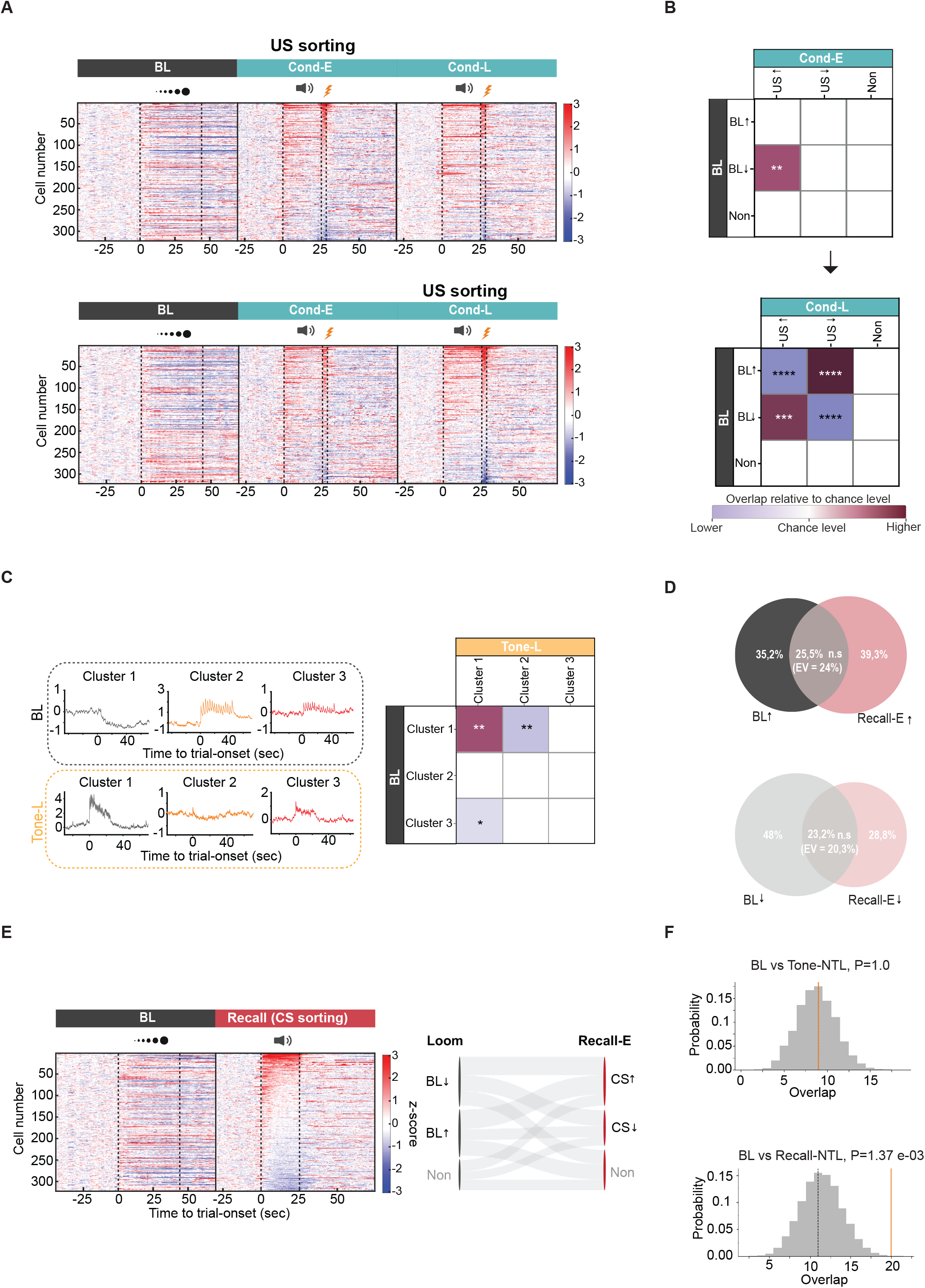
Distinct relationships between loom-activated neurons, US-activated neurons, and first-trial tone-responsive populations. (A) Heatmaps of single-cell z-scored activity during Loom, Cond-E, and Cond-L sessions, sorted by US response. Top, sorted by US response during Cond-E. Bottom, sorted by US response during Cond-L. Dotted vertical lines indicate CS/US onset and offset. (B) Contingency matrices between black loom-defined groups (BL↑, activated; BL↓, suppressed; Non, non-responsive) and US-defined groups (US↑, US↓, Non) during Cond-E (top) and Cond-L (bottom). Colors indicate direction of overlap (bordeaux, higher than chance; purple, lower than chance). Overlap significance vs. hypergeometric chance expectation; **p < 0.01, ***p < 0.001, ****p < 0.0001. (C) Left, PCA and k-mean clustering applied to BL- and Tone-L-activated cells (BL, k = 3, cluster 1=97 cells, cluster 2 = 46 cells, cluster 3=142 cells; Tone-L, k = 5, cluster 1=17 cells, cluster 2=173 cells, cluster 3=91 cells; clusters with minimal cell number are not shown. Right, Contingency matrix between BL and Tone-L clusters; overlap significance vs. hypergeometric chance expectation; *p < 0.05, **p < 0.01; asterisks denote greater-than-chance overlap (bordeaux) and significantly less overlap than expected (purple). (D) Comparison of the population response between the looming paradigm and recall of the conditioned tone. Venn diagrams showing overlap between loom-activated (BL↑, top) or loom-suppressed (BL↓, bottom) cells and cells activated (Recall-E↑) or suppressed (Recall-E↓) by the CS, respectively. Percentages indicate the proportion of overlapping cells; EV, overlap expected by chance. Overlap did not exceed chance in either comparison (n.s.). (E) Left, Heatmap of single-cell z-scored activity during Loom and Recall-E trials, sorted by the CS response. Right, Alluvial diagram tracking individual cells from BL-defined groups (BL↓, BL↑, Non) to CS-defined response category during Recall-E (CS↑, CS↓, Non). (F) Overlap between loom-activated (BL↑) neurons and non-time-locked (NTL) tone-responsive populations, tested against a permutation-based null distribution. Gray histograms show the distribution of overlap expected by chance across shuffles; the dashed vertical line marks the mean of the null distribution and the orange vertical line indicates the observed overlap. Top, overlap between BL↑ and non-time-locked tone-responsive cells during the Tone session (Tone-NTL) did not differ from chance—the observed overlap coincides with the null mean (P = 1.0). Bottom, overlap between BL↑ and non-time-locked recall tone-responsive cells (Recall-NTL) was significantly greater than chance, with the observed overlap falling in the upper tail of the null distribution (P = 1.37 × 10^−3^). P-values from permutation test.

**Supplementary Figure 3.**
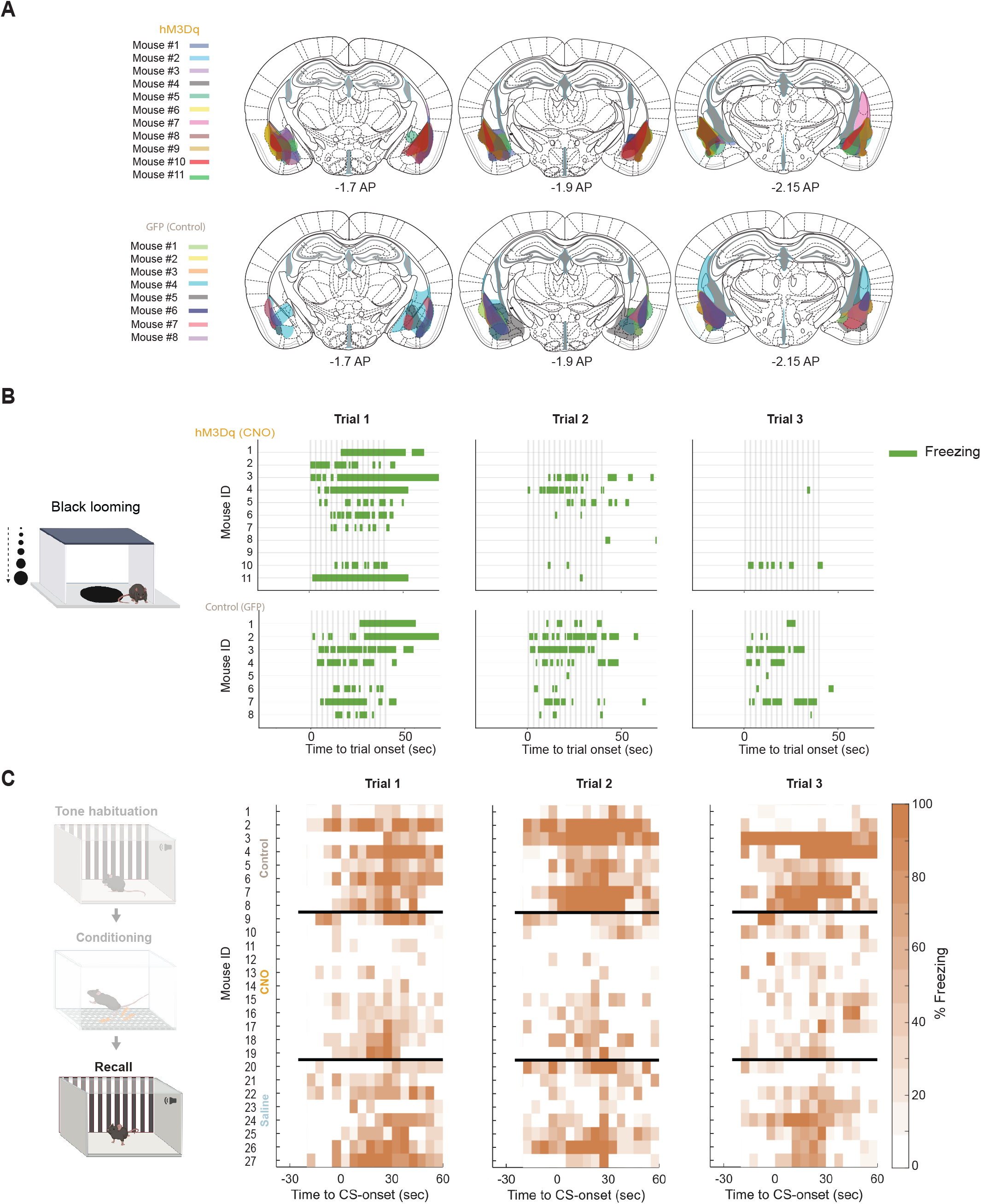
Viral expression and per-animal defensive behavior in the chemogenetic experiment. (A) Extent of viral expression for hM3Dq and control (GFP) mice across antero-posterior levels (approximately –1.7, –1.9 and –2.15 mm from bregma). (B) Per-animal time spent in freezing across trials in response to the black-looming stimulus, for hM3Dq versus control mice. (C) Per-animal freezing (% over time) to the CS at recall, for control (GFP) mice and for hM3Dq mice with CNO and with saline

**Supplementary Figure 4.**
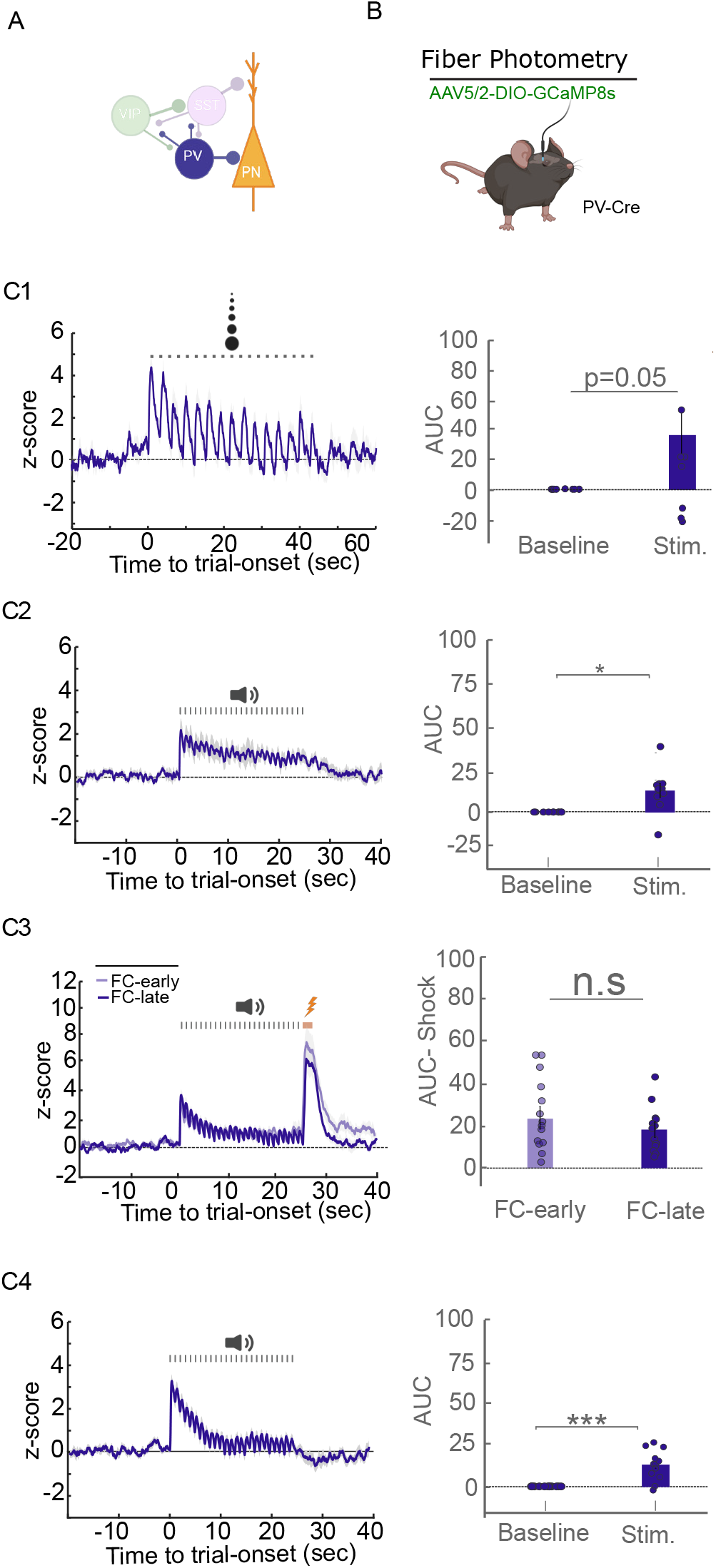
PV interneuron dynamics across learned and innate threats. (A) LA microcircuit schematic highlighting PV interneurons. (B) Fiber-photometry strategy: Cre-dependent jGCaMP8s (AAV5/2) in PV-Cre mice with an optic fiber over the LA (n = 10 mice). (C) Mean z-scored PV activity and corresponding area-under-the-curve (AUC) quantification (baseline vs. stimulus, paired t-test) for: (C1) the looming stimulus (p= 0.05); (C2) the neutral tone (p < 0.05); (C3) the US across early versus late conditioning trials (n.s.); and (C4) the CS during recall (p < 0.001). Error bars represent s.e.m.

**Supplementary Figure 5.**
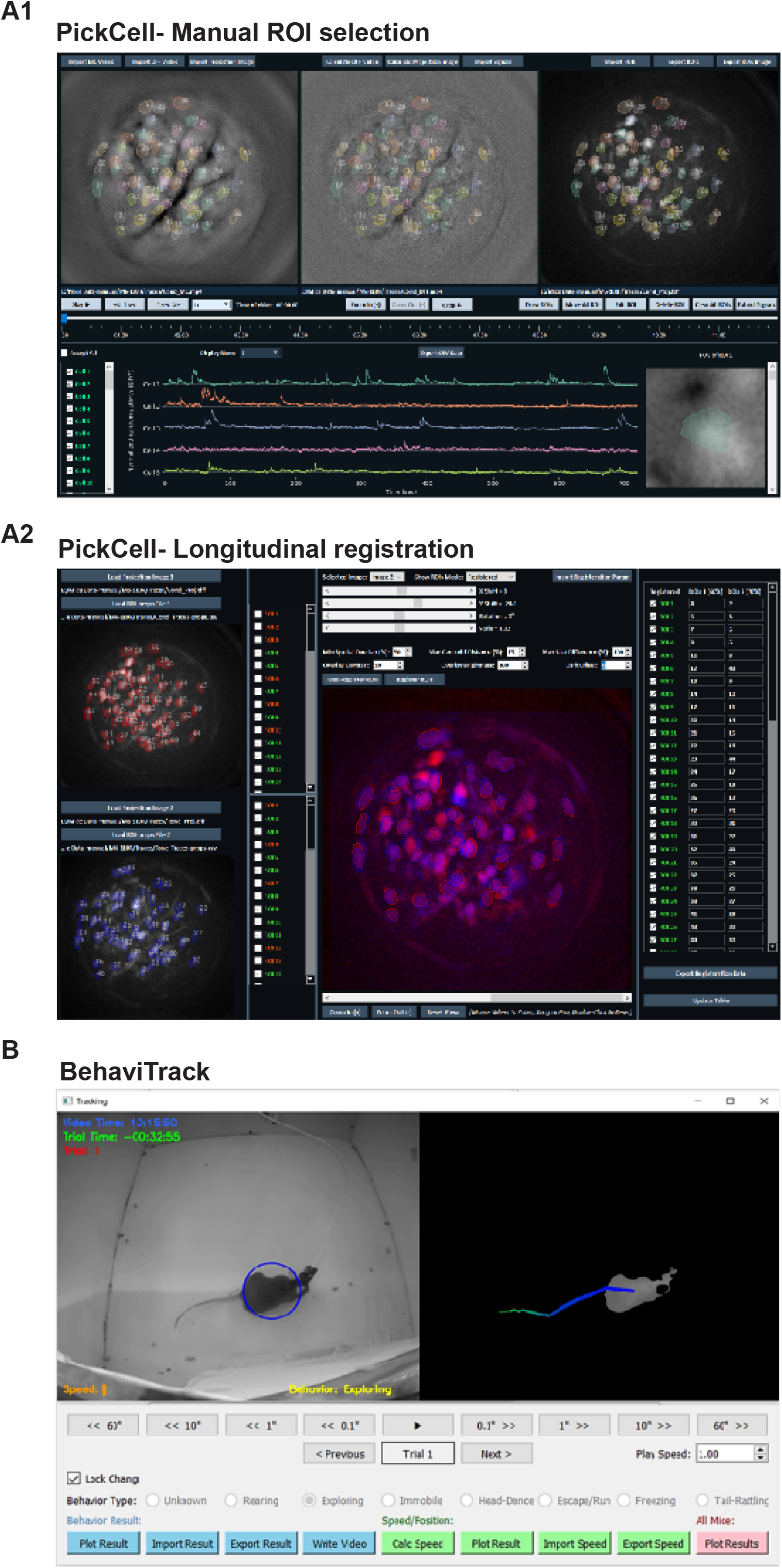
PickCell (manual ROI selection and longitudinal cell registration) and BehaviTrack (manual behavior scoring). (A1) Interface for manual ROI selection in PickCell. ROIs were manually drawn on motion-corrected videos, ΔF/F videos, or maximum projection images. The tool provides synchronized multi-panel views, zooming for precise ROI definition, and interactive ROI editing, including reshaping, moving, and deleting ROIs. ROI sets can be saved and reused across sessions. (A2) Longitudinal registration of cells across sessions. ROI maps from different sessions were aligned through rotation, scaling, and horizontal and vertical shifts to maximize spatial correspondence. PickCell supports overlay visualization, automated matching based on spatial overlap or centroid distance, and manual verification through an interactive registration table. (B) BehaviTrack is a custom Python-based tool for manual annotation of mouse behavior. It allows users to navigate videos at high temporal resolution and label behaviors. Key features include flexible video playback, visualization of average behavioral responses across animals, CSV import/export, automatic tracking of the animal’s position and speed within the arena, and camera angle correction when the recording setup is not perfectly aligned.

## References

1. Rosen, J. B. The Neurobiology of Conditioned and Unconditioned Fear: A Neurobehavioral System Analysis of the Amygdala. Behavioral and Cognitive Neuroscience Reviews (2004) doi:10.1177/1534582304265945.

2. Gross, C. T. & Canteras, N. S. The many paths to fear. Nature Reviews Neuroscience 13, 651–658 (2012).

3. Silva, B. A. et al. Independent hypothalamic circuits for social and predator fear. Nat Neurosci 16, 1731–1733 (2013).

4. Silva, B. A., Gross, C. T. & Gräff, J. The neural circuits of innate fear: detection, integration, action, and memorization. Learn Mem 23, 544–555 (2016).

5. Canteras, N. S. The medial hypothalamic defensive system: hodological organization and functional implications. Pharmacol Biochem Behav 71, 481–491 (2002).

6. Kunwar, P. S. et al. Ventromedial hypothalamic neurons control a defensive emotion state. Elife 4, (2015).

7. Kennedy, A. et al. Stimulus-specific hypothalamic encoding of a persistent defensive state. Nature 586, 730–734 (2020).

8. Li, C.-I., Maglinao, T. L. & Takahashi, L. K. Medial amygdala modulation of predator odor-induced unconditioned fear in the rat. Behav Neurosci 118, 324–332 (2004).

9. Johansen, J. P., Cain, C. K., Ostroff, L. E. & LeDoux, J. E. Molecular mechanisms of fear learning and memory. Cell 147, 509–524 (2011).

10. Johansen, J. P. et al. Hebbian and neuromodulatory mechanisms interact to trigger associative memory formation. Proc Natl Acad Sci U S A 111, E5584–92 (2014).

11. Quirk, G. J., Repa, C. & LeDoux, J. E. Fear conditioning enhances short-latency auditory responses of lateral amygdala neurons: parallel recordings in the freely behaving rat. Neuron 15, 1029–1039 (1995).

12. Rogan, M. T., Stäubli, U. V. & LeDoux, J. E. Fear conditioning induces associative long-term potentiation in the amygdala. Nature 390, 604–607 (1997).

13. Rumpel, S., LeDoux, J., Zador, A. & Malinow, R. Postsynaptic receptor trafficking underlying a form of associative learning. Science 308, 83–88 (2005).

14. Richardson, M. P., Strange, B. A. & Dolan, R. J. Encoding of emotional memories depends on amygdala and hippocampus and their interactions. Nat Neurosci 7, 278–285 (2004).

15. Maren, S. Synaptic mechanisms of associative memory in the amygdala. Neuron 47, 783–786 (2005).

16. Sah, P., Westbrook, R. F. & Lüthi, A. Fear conditioning and long-term potentiation in the amygdala: what really is the connection? Ann N Y Acad Sci 1129, 88–95 (2008).

17. Tovote, P., Fadok, J. P. & Lüthi, A. Neuronal circuits for fear and anxiety. Nat Rev Neurosci 16, 317–331 (2015).

18. Xu, C. et al. Distinct Hippocampal Pathways Mediate Dissociable Roles of Context in Memory Retrieval. Cell 167, 961–972.e16 (2016).

19. Herry, C. & Johansen, J. P. Encoding of fear learning and memory in distributed neuronal circuits. Nat Neurosci 17, 1644–1654 (2014).

20. Barsy, B. et al. Associative and plastic thalamic signaling to the lateral amygdala controls fear behavior. Nat Neurosci 23, 625–637 (2020).

21. Edeline, J. M. & Weinberger, N. M. Associative retuning in the thalamic source of input to the amygdala and auditory cortex: receptive field plasticity in the medial division of the medial geniculate body. Behav Neurosci 106, 81–105 (1992).

22. Maren, S. & Quirk, G. J. Neuronal signalling of fear memory. Nat Rev Neurosci 5, 844–852 (2004).

23. Bindi, R. P., Baldo, M. V. C. & Canteras, N. S. Roles of the anterior basolateral amygdalar nucleus during exposure to a live predator and to a predator-associated context. Behav Brain Res 342, 51–56 (2018).

24. Martinez, R. C., Carvalho-Netto, E. F., Ribeiro-Barbosa, E. R., Baldo, M. V. C. & Canteras, N. S. Amygdalar roles during exposure to a live predator and to a predator-associated context. Neuroscience 172, 314–328 (2011).

25. Kang, S. J. et al. A central alarm system that gates multi-sensory innate threat cues to the amygdala. Cell Rep 40, 111222 (2022).

26. Khalil, V. et al. Subcortico-amygdala pathway processes innate and learned threats. Elife 12, (2023).

27. Yilmaz, M. & Meister, M. Rapid innate defensive responses of mice to looming visual stimuli. Curr Biol 23, 2011–2015 (2013).

28. Dunn, T. W. et al. Neural Circuits Underlying Visually Evoked Escapes in Larval Zebrafish. Neuron 89, 613–628 (2016).

29. Fotowat, H. & Engert, F. Neural circuits underlying habituation of visually evoked escape behaviors in larval zebrafish. Elife 12, (2023).

30. Shang, C. et al. Divergent midbrain circuits orchestrate escape and freezing responses to looming stimuli in mice. Nat Commun 9, 1232 (2018).

31. Evans, D. A. et al. A synaptic threshold mechanism for computing escape decisions. Nature 558, 590–594 (2018).

32. Lecca, S. et al. Heterogeneous Habenular Neuronal Ensembles during Selection of Defensive Behaviors. Cell Rep 31, 107752 (2020).

33. Zhang, Y. et al. Fast and sensitive GCaMP calcium indicators for imaging neural populations. Nature 615, 884–891 (2023).

34. Ziv, Y. et al. Long-term dynamics of CA1 hippocampal place codes. Nat Neurosci 16, 264–266 (2013).

35. Grewe, B. F. et al. Neural ensemble dynamics underlying a long-term associative memory. Nature 543, 670–675 (2017).

36. Gründemann, J. et al. Amygdala ensembles encode behavioral states. Science 364, (2019).

37. Blair, H. T., Schafe, G. E., Bauer, E. P., Rodrigues, S. M. & LeDoux, J. E. Synaptic plasticity in the lateral amygdala: a cellular hypothesis of fear conditioning. Learn Mem 8, 229–242 (2001).

38. Pape, H.-C. & Pare, D. Plastic synaptic networks of the amygdala for the acquisition, expression, and extinction of conditioned fear. Physiol Rev 90, 419–463 (2010).

39. Salay, L. D., Ishiko, N. & Huberman, A. D. A midline thalamic circuit determines reactions to visual threat. Nature 557, 183–189 (2018).

40. Wolff, S. B. E. et al. Amygdala interneuron subtypes control fear learning through disinhibition. Nature 509, 453–458 (2014).

41. Letzkus, J. J., Wolff, S. B. E. & Lüthi, A. Disinhibition, a Circuit Mechanism for Associative Learning and Memory. Neuron 88, 264–276 (2015).

42. Fadok, J. P. et al. A competitive inhibitory circuit for selection of active and passive fear responses. Nature 542, 96–100 (2017).

43. Krabbe, S., Gründemann, J. & Lüthi, A. Amygdala Inhibitory Circuits Regulate Associative Fear Conditioning. Biol Psychiatry 83, 800–809 (2018).

44. Krabbe, S. et al. Adaptive disinhibitory gating by VIP interneurons permits associative learning. Nat Neurosci 22, 1834–1843 (2019).

45. Favila, N. et al. Heterogeneous plasticity of amygdala interneurons in associative learning and extinction. Nat Commun 16, 9926 (2025).

46. Armbruster, B. N., Li, X., Pausch, M. H., Herlitze, S. & Roth, B. L. Evolving the lock to fit the key to create a family of G protein-coupled receptors potently activated by an inert ligand. Proc Natl Acad Sci U S A 104, 5163–5168 (2007).

47. Feng, J. et al. Monitoring norepinephrine release in vivo using next-generation GRAB sensors. Neuron 112, 1930–1942.e6 (2024).

48. Romanski, L. M. & LeDoux, J. E. Equipotentiality of thalamo-amygdala and thalamo-cortico-amygdala circuits in auditory fear conditioning. J Neurosci 12, 4501–4509 (1992).

49. Shang, C. et al. A parvalbumin-positive excitatory visual pathway to trigger fear responses in mice. Science 348, 1472–1477 (2015).

50. Stefanelli, T., Bertollini, C., Lüscher, C., Muller, D. & Mendez, P. Hippocampal Somatostatin Interneurons Control the Size of Neuronal Memory Ensembles. Neuron 89, 1074–1085 (2016).

51. Morrison, D. J. et al. Parvalbumin interneurons constrain the size of the lateral amygdala engram. Neurobiol Learn Mem 135, 91–99 (2016).

52. Hayley, S., Borowski, T., Merali, Z. & Anisman, H. Central monoamine activity in genetically distinct strains of mice following a psychogenic stressor: effects of predator exposure. Brain Res 892, 293–300 (2001).

53. Hu, H. et al. Emotion enhances learning via norepinephrine regulation of AMPA-receptor trafficking. Cell 131, 160–173 (2007).

54. Do Monte, F. H. M., Canteras, N. S., Fernandes, D., Assreuy, J. & Carobrez, A. P. New perspectives on beta-adrenergic mediation of innate and learned fear responses to predator odor. J Neurosci 28, 13296–13302 (2008).

55. Liu, Y. et al. A single fear-inducing stimulus induces a transcription-dependent switch in synaptic AMPAR phenotype. Nat Neurosci 13, 223–231 (2010).

56. Tan, B. Z. et al. A neuromodulatory circuit-to-molecular pathway for reformatting aversive memories during recall. Neuron 114, 1848–1862.e7 (2026).

57. Faber, E. S. L. et al. Modulation of SK channel trafficking by beta adrenoceptors enhances excitatory synaptic transmission and plasticity in the amygdala. J Neurosci 28, 10803–10813 (2008).

58. Tully, K., Li, Y., Tsvetkov, E. & Bolshakov, V. Y. Norepinephrine enables the induction of associative long-term potentiation at thalamo-amygdala synapses. Proc Natl Acad Sci U S A 104, 14146–14150 (2007).

59. Zador, A. M. A critique of pure learning and what artificial neural networks can learn from animal brains. Nat Commun 10, 3770 (2019).

60. Mondoloni, S., Mameli, M. & Congiu, M. Reward and aversion encoding in the lateral habenula for innate and learned behaviours. Transl Psychiatry 12, 3 (2022).

61. Barbano, M. F. et al. VTA Glutamatergic Neurons Mediate Innate Defensive Behaviors. Neuron 107, 368–382.e8 (2020).

